# Models of SIV rebound after treatment interruption that involve multiple reactivation events

**DOI:** 10.1101/2020.07.28.221226

**Authors:** Christiaan H. van Dorp, Jessica M. Conway, James B. Whitney, Dan H. Barouch, Alan S. Perelson

## Abstract

In order to assess the efficacy of novel HIV-1 treatments leading to a functional cure, the time to viral rebound is frequently used as a surrogate endpoint. The longer the time to viral rebound, the more efficacious the therapy. In support of such an approach, mathematical models serve as a connection between the size of the latent reservoir and the time to HIV-1 rebound after treatment interruption. The simplest of such models assumes that a single successful latent cell reactivation event leads to observable viremia after a period of exponential viral growth. Here we consider a generalization developed by Pinkevych *et al.* and Hill *et al.* of this simple model in which multiple reactivation events can occur, each contributing to the exponential growth of the viral load. We formalize and improve the previous derivation of the dynamics predicted by this model, and use the model to estimate relevant biological parameters from SIV rebound data. We confirm a previously described effect of very early antiretroviral therapy (ART) initiation on the rate of recrudescence and the viral load growth rate after treatment interruption. We find that every day ART initiation is delayed results in a 39% increase in the recrudescence rate, and a 11% decrease of the viral growth rate. We show that when viral rebound occurs early relative to the viral load doubling time, a model with multiple successful reactivation events fits the data better than a model with only a single successful reactivation event.

**Author Summary:** HIV-1 persists during suppressive antiretroviral therapy (ART) due to a reservoir of latently infected cells. When ART is stopped, HIV generally rebounds within a few weeks. However, there is a small fraction of patients that do not rebound over a period of months or years. A variety of treatments are being tested for their ability to reduce the size of the latent reservoir, to induce effective immune responses against the virus, or to prevent or prolong the time to viral rebound after ART interruption. These novel treatments are typically first tested in SIV infected macaques, and the efficacy of the treatment assessed by interrupting ART and measuring the time to viral rebound. Here, we develop and test a mathematical and statistical model that describes the process of viral rebound. The model can be used for statistical inference of the efficacy of newly developed treatments. Importantly, the model takes into account that multiple recrudescence events can precede rebound. We test the model using data from early treated SIV infected macaques.

## Introduction

HIV and SIV are able to persist despite antiretroviral therapy (ART) because of a long-lived reservoir of latently infected CD4^+^ T cells [26]. Recent studies have shown that the latent reservoir is established very early after infection [43, 29, 8], and that the seeding of the reservoir can only be prevented when ART starts extremely early [44]. Other studies have focused on the effect of potentially curative treatment strategies that might extend remission after interruption of ART [5, 3, 4].

In all these studies an important observable is the time between treatment interruption and viral rebound, i.e. the first time the viral load (VL) becomes observable. Under the common assumption that rebound results from reactivation of latently infected cells [cf. 18, 31, 10], and that the rate at which the latent population reactivates is proportional to the size of the latent reservoir, the time to viral rebound can be used to gauge the reservoir size. Some curative strategies aim to reduce the size of the reservoir by administering latency reversing agents such as vorinostat [2], romidepsin [40], and TLR7 agonists [4]. The time to rebound can then we used as an indication of the effectiveness of the treatment, consistent with the aforementioned assumption [17, 18, 35].

The simplest model of rebound combines an exponentially distributed waiting time for a recrudescence event with subsequent exponential growth of the VL (henceforth, this is referred to as the “single-reactivation model”). Such a model has been used to estimate the reactivation rate of cells from the reservoir in HIV-1 patients undergoing ART interruption [31]. The main conclusion of this study— reactivation occurs on average every 5-8 days—resulted in some discussion about the sensitivity of the aforementioned result to inter-patient variability of the model parameters [19, 34]. From this discussion, an interesting and slightly more complex model of viral rebound emerged [19, 34] that takes into account the possibility that multiple latently infected cells reactivate within a short time interval, and that each of these reactivation events contributes to VL growth (we hereafter refer to this model as the “multiple-reactivation model”). We refer to a reactivation event that leads to an exponentially growing and potentially observable lineage of actively infected cells as a “successful reactivation event” or “recrudescence event”, since by chance alone a reactivation can also lead to a sequence of infection events that ultimately goes extinct [17, 10, 30, 9].

The occurrence of multiple recrudescence events is not merely a theoretical hypothesis, but has recently been observed *in vivo*. In one study, phylogenetic analysis has revealed that HIV-1 rebound is seeded from multiple anatomical sites [11]. In another study, treatment interruption experiments with macaques infected with a genetically barcoded SIV strain showed that many cells successfully reactivate from the latent reservoir [13]. In the latter study, the multiple-reactivation model was used to analyze the viral rebound data [13, 33], underpinning the current interest in this model. Moreover, in a recent analysis of potentially curative treatment effects the multiple-reactivation model was used as a bridge between stochastic and deterministic reactivation domains [35]. Here we present an improvement of the multiple-reactivation model that we derive using a Poisson counting process. Although the average behavior of our improved model is only marginally different from the previous version, our approach allows us to not only model the expected viral load rebound curve, but also the deviation from this expectation. Most importantly, this enables us to derive a parametric expression for the distribution of the time-to-rebound, that can be used for parameter inference from rebound data.

We test the improved model using data from SIV infected macaques that are put on ART at different times post infection and exhibit varying viral rebound dynamics [43, 44]. We find very strong statistical evidence in favor of the multiple-reactivation model over the single-reactivation model. We attribute this superior model performance to the fact that it better explains the data from macaques that rebound soon after ART cessation and exhibit relatively slow exponential growth of the VL. We argue that whenever such data is used for inference about the effects of experimental curative treatments in delaying viral rebound, the multiple-reactivation model should be used to estimate the relevant parameters. Our refined multiple-reactivation model fits the data only slightly better than the approximation developed earlier by Pinkevych *et al.* [34]. However, using an example, we show that our model can be generalized further to include more complex features of reservoir and rebound dynamics, such as heterogeneity of the reservoir in terms of clone-specific growth rates.

## Results

We start by mathematically defining the multiple-reactivation model and deriving the mean behavior and deviation from the mean of this model. We then use these quantities to derive an approximate probability distribution of the time to viral rebound, and assess whether this approximation is reliable. This time-to-rebound distribution is then used to infer the rate of recrudescence from a heterogeneous set of SIV rebound data. This inference allows us to quantify the effect of ART initiation time on the recrudescence rate and viral growth rate, and to compare our multiple-reactivation model with the simpler single-reactivation model. We identify two mechanisms that make the multiple-reactivation model better suited for modeling rebound data than the single-reactivation model. Finally, using simulated data sets, we test how sensitive the model is to parameter and model misspecification.

### The multiple- and single-reactivation models

We start by constructing a model that predicts the short-term SIV or HIV viral dynamics following the cessation of ART including viral rebound to detectable viremia and subsequent exponential growth of the VL. In our modeling we rely on the common, central assumption that activation of latently infected cells drives viral rebound [18, 31, 10]. Specifically, we assume that the activation of a latently infected cell can be followed by viral production, which in turn may lead to infection of additional cells. Viral rebound is caused by exponential growth in resultant viral lineages. We refer to a latent cell reactivation that leads to exponential growth as a “successful reactivation event” or a “recrudescence event”, to explicitly make the distinction with reactivation events leading to a viral lineage that by chance goes extinct while the population size is still small. We provide an overview of the models we employ in this study with full details provided in the Methods. A synopsis of the parameters and variables used is given in Table 1.

**Table 1:** An overview of the parameters and variables. See also Table 2 for additional parameters of the Bayesian mixed-effects model.

| symbol | unit | description |
| --- | --- | --- |
| $N_t$ | - | number of recrudescence events at time $t$ post treatment interruption |
| $V_t$ | copies mL<sup>-1</sup> | VL at time $t$ post treatment interruption |
| $T_i$ | d | time of the $i$-th successful reactivation event. |
| $g$ | d<sup>-1</sup> | exponential growth rate of the VL |
| $v_0$ | copies mL<sup>-1</sup> | average initial viral concentration caused by a single successful reactivation |
| $\lambda$ | d<sup>-1</sup> | recrudescence rate, the number of latently infected cells that successfully reactivate per day. |
| $K(\theta)$ | - | cumulant-generating function of the viral load $V_t$. |
| $\kappa_1$ | copies mL<sup>-1</sup> | first cumulant, equal to the expectation of the VL ($\mathbb{E}[V_t]$). |
| $\kappa_2$ | copies<sup>2</sup>mL<sup>-2</sup> | second cumulant, equal to the variance of the VL ($\text{Var}[V_t]$). |
| $\tilde{V}_t$ | copies mL<sup>-1</sup> | conditionally deterministic approximation of the viral load process $V_t$. |
| $\ell$ | copies mL<sup>-1</sup> | limit of detection of the VL assay. |
| $\tau$ | d | viral rebound time, satisfies the equation $V_\tau = \ell$ |
| $S(t)$ | - | fraction of subjects in remission at time $t$ post treatment interruption |
| $f(t; \lambda, g, v_0, \ell)$ | d<sup>-1</sup> | approximation of the probability density function of the rebound time $\tau$ |
| $\hat{T}_1$ | d | first recrudescence time extrapolated from the rebound time using the growth rate and the initial VL, assuming simple exponential growth. |
| $t_{\text{ART}}$ | d | days post infection that ART was initiated |
| $G_i$ | d<sup>-1</sup> | random, clone-specific exponential growth rate. |
| $\sigma_G$ | d<sup>-1</sup> | standard deviation of growth rates ($G_i$) of the clones comprising the SIV reservoir |

Mathematically, the multiple-reactivation model is a combination of a stochastic Poisson counting process *N*_*t*_ with rate or intensity *λ* and deterministic exponential viral growth *v*_0_*e*^*gt*^ with growth rate *g* and initial value *v*_0_. The Poisson process *N*_*t*_ counts the number of latently infected cells that have been reactivated and successfully establish a lineage of exponentially growing infected cells at time *t* after treatment interruption. Under the assumption that such successful reactivation events start occurring after therapy interruption at time *t* = 0, we have *N*_0_ = 0 and *N*_*t*_ ∼ Poisson(*λt*). The exponential curve *v*_0_*e*^*gt*^ describes the contribution to the total VL of such a lineage. The total VL at time *t* is the weighted sum of such exponential functions:

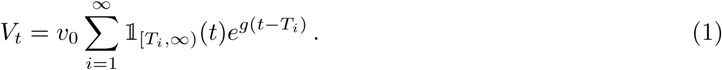

Here, the indicator function 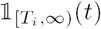 equals 1 if *t* ≥ *T*_*i*_ and 0 otherwise. The random times *T*_*i*_ are the jump times of the Poisson process, corresponding to the times that different latently infected cells successfully reactivate. An example realization of the random process *V*_*t*_ given by Eq 1 is shown in Fig 1A. Notice that there might be some delay between the moment of reactivation and successful reactivation. For instance, it might be possible that the reactivation of a latently infected cell happens before treatment interruption. The times *T*_*i*_ correspond to the moments that lineages initiated by reactivation become large enough, the meaning of which we explore in the Discussion.

**Figure 1:**
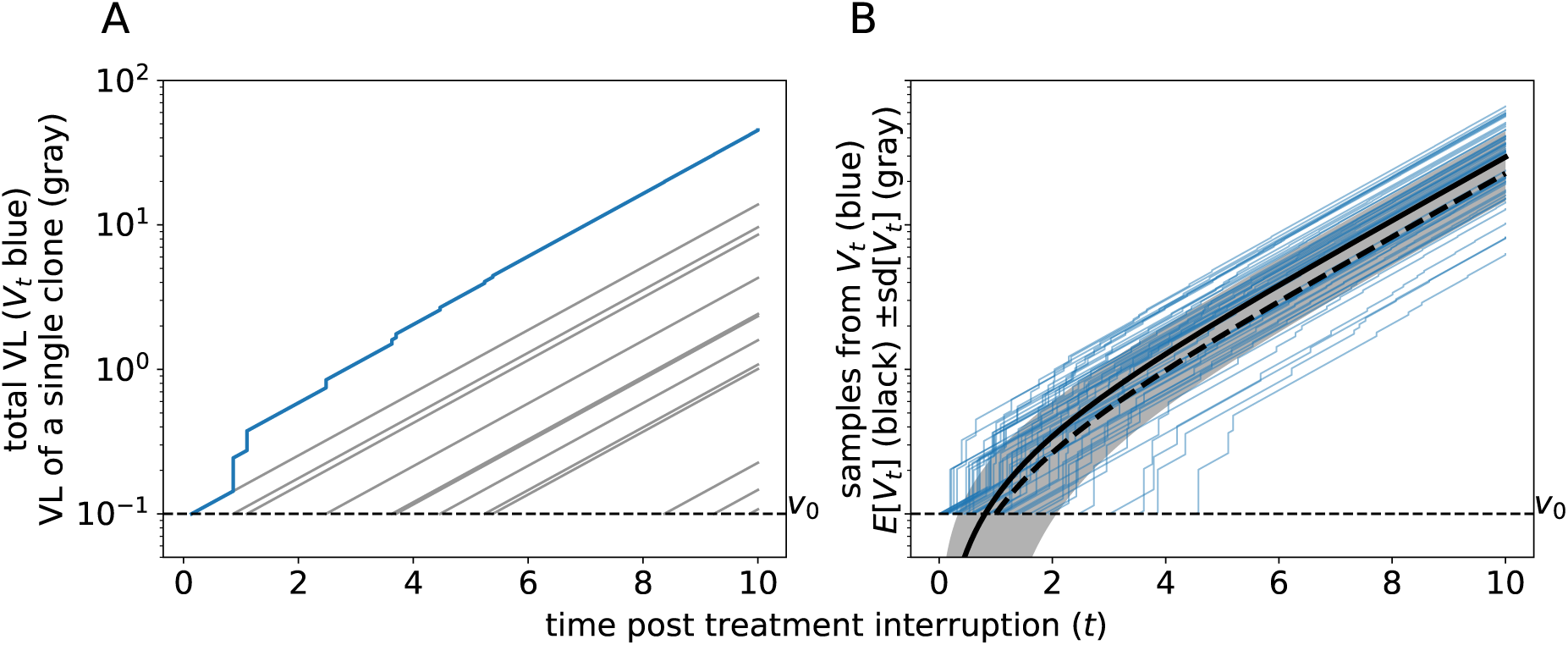
Simulations of the multiple-reactivation model. (A) Graphical representation of Eq 1. The gray lines indicate the exponential growth curves of individual clones that originated from a single successful reactivation from the latent reservoir. The blue curve represents the total VL, i.e. the sum of the gray lines. (B) Comparison between the expectation of the process *V*_*t*_ (in black) and realizations sampled from this process (in blue). The mean *±* standard deviation (sd) of *V*_*t*_ is shown as a gray band. The dashed thick curve corresponds to the approximation 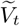 with *t*_0_ = 1*/λ* d. Parameters: *g* = 0.5 d^−1^, *λ* = 1.0 d^−1^, and *v*_0_ = 0.1 copies mL^−1^

In previous analyses of Eq 1 by Pinkevych *et al.* [34] and others [13, 33, 35], the dynamics of the process *V*_*t*_ after the initial reactivation event *T*_1_ = *t*_0_ was simplified using a deterministic approximation. The subsequent recrudescence times *T*_2_, *T*_3_, … were assumed to be exactly 1*/λ* days apart, which is the average time between two succeeding jumps of the Poisson process. With the aid of some further simplifications [see 34, 13, 35, or Methods], the following expression was obtained for the total VL at time *t* ≥ *t*_0_ after ART suspension:

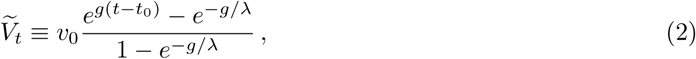

where the tilde over the *V* is used to indicate that this is an approximation. In the Methods section, we use the cumulant-generating function (CGF), together with some basic facts about the Poisson process to derive a functional form for the expectation of *V*_*t*_, where we no longer constrain recrudescence times to be 1*/λ* days apart:

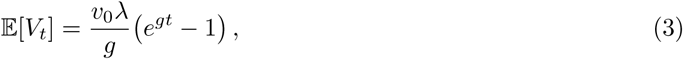

Moreover, the same CGF technique allows us to find all other cumulants (or moments) of the distribution of *V*_*t*_. For instance, we show the variance is given by

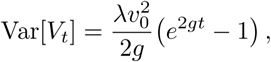

and the third cumulant, which has the same sign as the skewness, equals 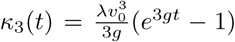. The expected trajectory of *V*_*t*_ and the standard deviation are shown in Fig 1B. To compare the difference between Eq 3 and Eq 2, the graph of 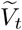 is shown as a thick dashed curve in Fig 1B. This example shows that the approximation 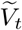 slightly under-estimates the expected VL (the thick black line in Fig 1B). However, the primary advantage of our improvement comes from the additional statistical properties of viral rebound dynamics that it allows us to compute, which is useful for the estimation of parameters such as the recrudescence rate *λ* (see below).

We term this model the “multiple-reactivation model” since viral load is modeled as the sum of viral lineages generated by multiple recrudescence events. In the following we contrast predictions from our refined multiple-reactivation model with the “single-reactivation model”, in which rebound VL is assumed to be associated with the viral lineage resulting from a single latent cell activation only [cf. 31]. Hence, for this single-reactivation model we consider only the first recrudescence event at time *T*_1_ ∼ Exponential(*λ*), and ignore the effects of any subsequent reactivations from the reservoir. Given that *T*_1_ = *t*_0_, we therefore get the following simple expression for *t* ≥ *t*_0_:

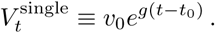

### The distribution of time-to-rebound

In treatment-interruption experiments, the main quantity of interest is the time-to-rebound, which we denote by *τ*. In order to properly infer the recrudescence rate *λ*—a proxy for replication competent reservoir size—from viral rebound data, a statistical model that expresses the likelihood of the time-to-rebound in terms of the model parameters is desirable. The predicted distributions of the time-to-rebound under the multiple-reactivation model and single-reactivation model naturally differ, because multiple reactivation events, early after treatment interruption, skew the time-to-rebound towards lower values. This means that because of these multiple recrudescence events prior to viral rebound, each of which causing a jump in the viral load, the growth of the still unobservable VL is faster than exponential growth at rate *g*. We refer to this as “early faster-than-exponential growth”. Using an exponential distribution for the first recrudescence time *T*_1_, and the approximation 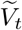 given in Eq 2, the rate of successful reactivation *λ* [34, 13], and the initial contribution *v*_0_ of such a reactivation event [33] have been estimated with likelihood-based methods. However, this conditionally deterministic approximation does not take the uncertainty due to secondary recrudescence events occurring at different intervals into account, a shortcoming which we fix with our fully stochastic model.

Using a diffusion approximation of the process *V*_*t*_ allows us to derive a convenient parametric form of the distribution of the time-to-rebound (given by Eq 6 in Methods). The time-to-rebound (*τ*) is defined as the first time the virus load crosses a threshold *ℓ* corresponding to the limit of detection (LoD; typically 50 RNA copies per mL) of the assay used to measure SIV or HIV RNA. Our parametric distribution depends on *ℓ* and the parameters *v*_0_, *λ*, and *g* and can be used to estimate these parameters directly from time-to-rebound data using methods such as maximum likelihood. In order to test if the diffusion approximation is justified, we simulated the process *V*_*t*_ and compared the empirical distribution of the time to rebound with the parametric approximation (see Fig 2). When successful reactivation is fast (*λ* ≥ 1 d^−1^), the simulations and our approximation are in excellent agreement (by visual inspection; Fig 2 top and middle panels).

**Figure 2:**
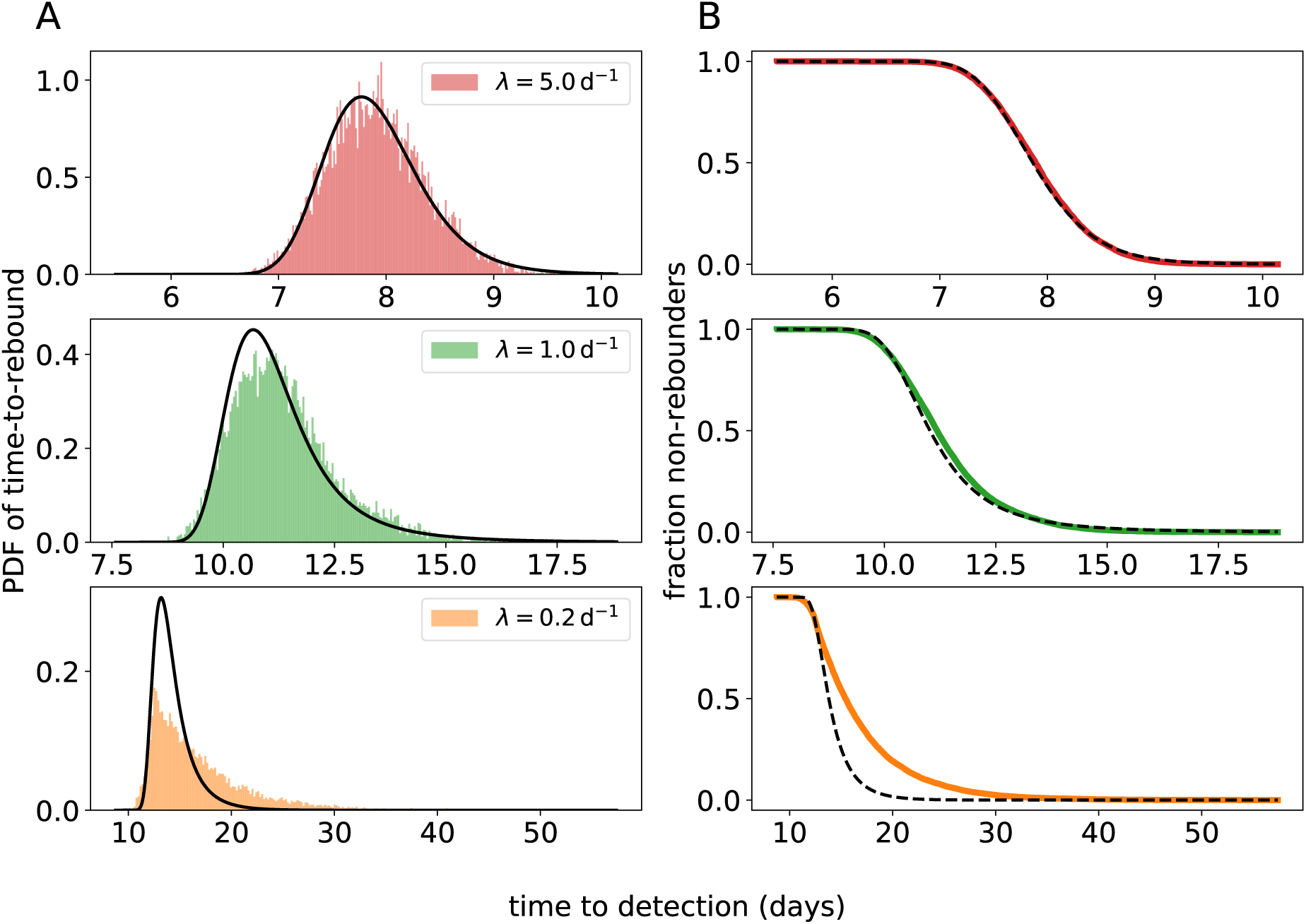
Comparison between the approximation for the time-to-rebound distribution and simulated rebound times. The simulated empirical distributions are shown in color, and our approximation is shown in black. (A) The probability density function (PDF; defined by Eq 6 and 7). (B) The survival function (i.e. the fraction of subjects *S*(*t*) that do not have a detectable VL at time *t*). For the top, middle, and bottom panels different values of *λ* are used (*λ* = 5 d^−1^, 1 d^−1^, and 0.2 d^−1^ respectively). Notice the different time scale on the *x*-axes. For the remaining parameters, we used the values: *g* = 0.5 d^−1^, *v*_0_ = 0.1 copies mL^−1^, LoD *ℓ* = 50 copies mL^−1^.

However, when the successful-reactivation rate is small (*λ* = 0.2 d^−1^), the diffusion approximation breaks down (Fig 2 bottom panels), as the time to rebound is mostly determined by the first successful reactivation, and hence by the exponentially distributed initial recrudescence time *T*_1_. Further, the distribution of the diffusion approximation of *V*_*t*_ at time *t* is a Gaussian 𝒩 (*κ*_1_(*t*), *κ*_2_(*t*)), which is symmetric. As *κ*_3_(*t*) > 0 for *t* > 0 the exact distribution of *V*_*t*_ is in fact right-skewed. When we fit the multiple-reactivation model to data below, we account for these discrepancies by explicitly modeling the first reactivation time *T*_1_ as a so-called latent variable of the statistical model. The diffusion approximation for the time-to-rebound distribution (Eq 6) is then used to model the difference *τ* − *T*_1_, and we set an initial condition 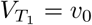. This ensures that the model can be used for inference irrespective of whether remission is short or long, and even with data sets containing heterogeneous rebound times, as we will demonstrate below.

In the S1 Text we explore two other approximations of the rebound-time distribution that behave better for small values of *λ*. First, we replaced the Gaussian distribution *𝒩* (*κ*_1_, *κ*_2_) with a Gamma distribution, for which we matched the mean and variance with *κ*_1_ and *κ*_2_, respectively. Like the distribution of *V*_*t*_, the Gamma distribution has a positive skewness, which results in a greater similarity between the approximate rebound-time distribution and simulations when the recrudescence rate is small (Fig S3). Second, instead of diffusion, we applied the so-called WKB approximation to the process *V*_*t*_, which gave even better results for small recrudescence rates than the Gamma-law approximation (Fig S4). Unfortunately, both these improved approximations are more difficult to implement in standard parameter-inference frameworks. For this practical reason, we use the more tractable diffusion approximation in our data analysis below.

### Analysis of SIV rebound data

To assess the performance of the multiple-reactivation model and our diffusion approximation with respect to actual data, we employ the results of treatment interruption experiments with the macaque SIV model [43, 44]. This data set consists of longitudinal VL measurements from macaques for whom treatment was initiated early and at varying time points after SIV challenge in different groups of animals. The time of ART initiation has been shown to be a predictor for the time-to-rebound with early SIV treatment leading to delayed rebound [43]. Moreover, in the same study [43] it was found that the rate of exponential growth of the VL after viral rebound is decreased when ART is initiated later, perhaps because immune responses develop due to higher antigen concentrations. Hence, the data set contains SIV rebound time series with varying exponential growth rates and rebound times.

Of the 36 macaques in the data set, *n* = 25 showed viral rebound during the 16 week observation period after treatment interruption. As VL is measured at discrete times, the actual time of viral rebound (*τ*) has to be interpolated from these VL measurements. In addition, estimating the recrudescence rate (*λ*) requires that we also estimate the viral growth rate (*g*), and since we expect *g* and *λ* to be correlated, additional data that informs the growth rate helps to estimate both *g* and *λ* more accurately. In order to infer *τ* and estimate *g* from the VL time series for each macaque, we fit a logistic growth model [cf. 43] to the initial VL data points of the time series. We manually selected time points that are consistent with logistic growth (the gray data points in Fig 3A and S1 were excluded). We opted for logistic instead of exponential growth because fitting an exponential growth model to non-linear rebound data (Fig 3A and S1) can result in an under-estimation of the growth rate [35]. Because for many macaques the number of observations that can inform these estimates is limited, we used a mixed effects model to estimate the growth rate *g*, using the time of ART initiation (*t*_ART_) as a covariate. Similarly, *t*_ART_ is used as a covariate for estimating the reactivation rate *λ*, which again has a random effect for each macaque. The statistical model is defined in full detail in the Methods.

**Figure 3:**
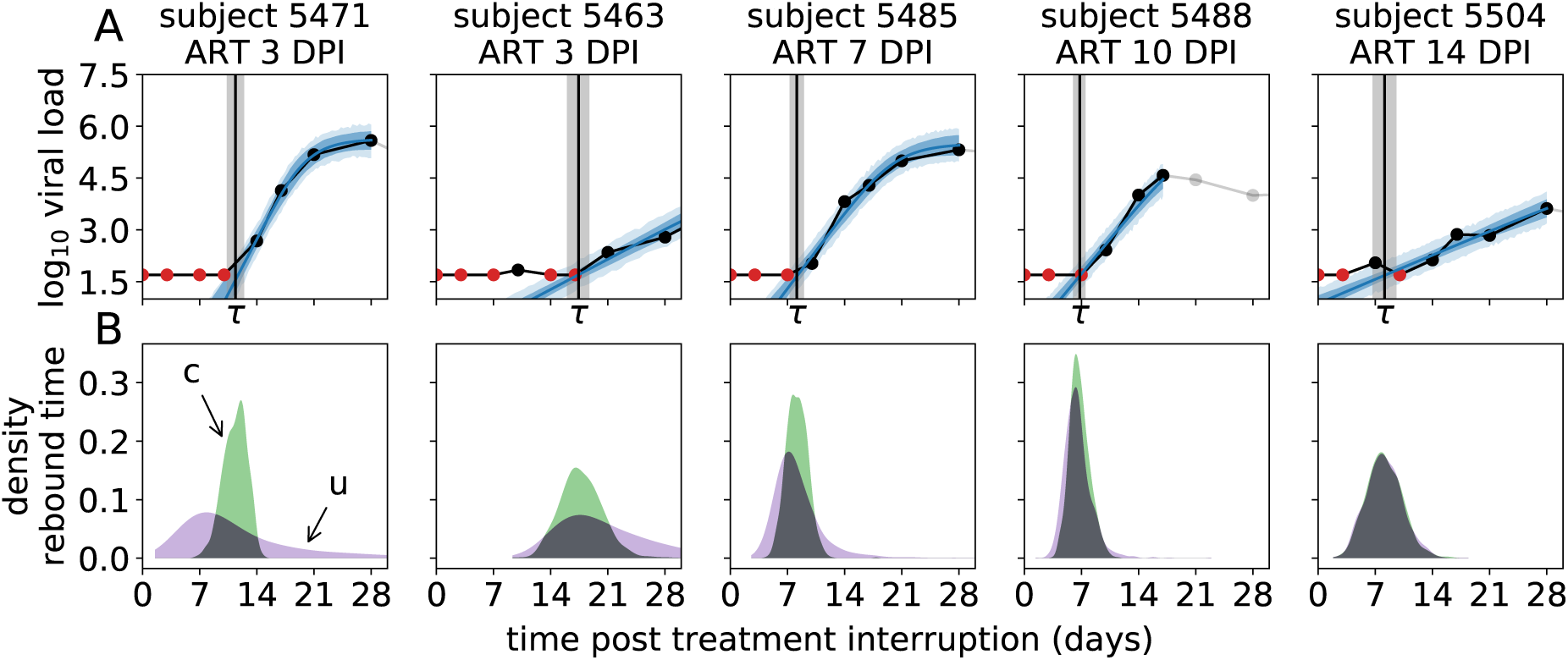
Representative examples of the fits of the mixed-effects model to the VL rebound time series. (A) The top panels (DPI: days post infection) show the VL data (black dots connected by black lines, with red dots for left-censored observations; the grey dots are ignored) taken from macaques where ART was started at different days post infection (DPI), and the model prediction (blue lines: posterior mean; dark blue band: 50% credibility interval (CrI), light blue band: 50% posterior predictive interval). The estimated time-to-rebound (*τ*) is given by the vertical black line (gray band: 50% CrI). (B) The bottom panels show posterior predictive distribution of the time-to-rebound. The green distributions (c) are conditioned on the estimated time of the initial reactivation event, the purple distributions (u) are unconditional. Model fits and posterior predictive distributions for all 25 macaques are shown in Fig S1.

Following Pinkevych *et al.* [33], we fit the logistic growth model to the VL data in a Bayesian framework using MCMC (Fig 3A and S1). This way we are able to naturally factor the rebound time distribution for the multiple-reactivation model (Eq 6) into the likelihood. We have to fit the model to the data from all 25 macaques simultaneously, as it contains fixed and random effects. We write *α*_*λ*_ (and *α*_*g*_) for the fixed effect of *t*_ART_ on *λ* (and *g*, respectively; see Eq 9 in Methods). In accordance with previous analyses [43], we find that the time of ART initiation is a strong predictor of both the rate of reactivation and the exponential growth rate after rebound. The posterior probability of the fixed effect *α*_*λ*_ of the time of ART initiation on the recrudescence rate ℙ[*α*_*λ*_ < 0] < 10^−3^ strongly suggests that *α*_*λ*_ > 0, i.e. that rebound occurs more rapidly when ART is initiated later. For the fixed effect *α*_*g*_ of the time of ART initiation of the viral rebound growth rate we find the posterior probability ℙ[*α*_*g*_ > 0] = 0.002, suggesting that it is highly likely that the growth rate will slow down with later ART initiation. Here the statistical significance of the effect of treatment initiation time is much larger than found previously [43]. This increased significance is due the inclusion of data from additional macaques [44], as exclusion of this data gave a posterior probability ℙ[*α*_*g*_ > 0] = 0.36.

The estimates of *λ* and *g* for individual macaques are shown in Fig 4A and B as a function of ART initiation time, and also listed in Table S1. The recrudescence rate is clearly influenced by the ART initiation time. We predict that each day ART is delayed, the recrudescence rate is increased by 39% (50% CrI: [30%, 46%]). Even though we find that the time ART starts is a significant predictor for the growth rate *g* after rebound, the standard deviation of the growth rate’s random effects (*σ*_*g*_) is about 5 times larger than that of the reactivation rate (*σ*_*λ*_; see Table S1). Nonetheless, we can estimate that each day ART is delayed, the growth rate decreases by 11% (50% CrI: [9%, 14%]).

**Figure 4:**
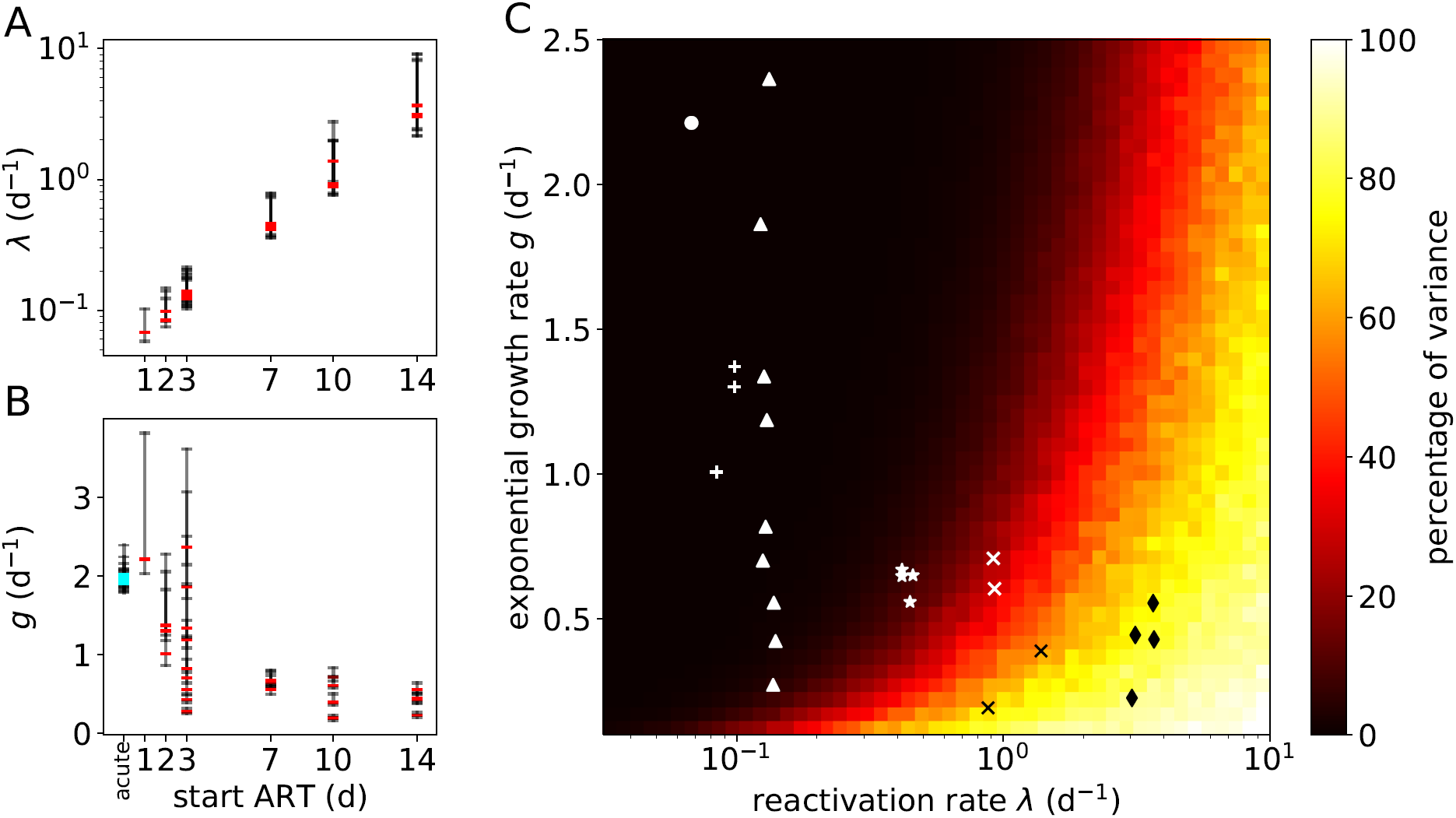
Estimates of recrudescence and growth rates from the SIV rebound data and the percentage of the variance of the time-to-rebound. (A) Point estimates (posterior modes; red) and 50% CrIs (black) of *λ* for each macaque as a function of the time ART was initiated. (B) Estimates of *g*. The cyan markers denote estimates of the growth rate for acute infections of 13 of the 25 macaques. These acute VL growth rates cluster around 2d^−1^. (C) Proportion of total variance due to secondary reactivation events. The heat map shows Var[*τ*_1_]*/*Var[*τ*_0_] · 100%, where *τ*_*i*_ := inf{*t* : *V*_*t*_ ≥ *R, V*_0_ = *i · v*_0_} is the rebound time (*i* = 0) or the time between the first successful reactivation and rebound (*i* = 1). Additional parameters are *v*_0_ = 0.1 copies mL^−1^ and *R* = 50 copies mL^−1^. The markers indicate the estimates from macaque SIV rebound experiments in which the macaques were treated, starting *t*_ART_ days after infection, with *t*_ART_ equal to 1 day (•), 2 days (**+**), 3 days (A), 7 days (*), 10 days (**×**), or 14 days (♦).

The latter observation is remarkably consistent with acute VL dynamics. As none of the macaques that were treated on day 0 (6h post-infection) showed viral rebound after ART cessation [44], we could not directly estimate the viral growth rates for these animals. However, we could compare the estimated growth rates after viral rebound with growth rates in the acute phase. Using again a simple random-effects logistic growth model, we were able to estimate viral growth rates during acute infection for the 13 out of the 25 macaques that showed observable viremia prior to ART initiation. These estimates are added to Fig 4B (cyan markers, located at “acute”). Our estimates for the acute growth rates are slightly higher than reported previously [43], possibly due to the use of a logistic growth model (see Methods). Using our estimates from the rebound data of the population-level growth rate (*µ*_*g*_, see Table 2) and fixed effect of *t*_ART_ (*α*_*g*_), we extrapolated the population-level growth rate for subjects treated at day 0 (using Eq 9). Our estimate of the population-level growth rate for the acute infection (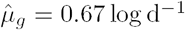; Table S1) falls within the 50% CrI of the extrapolated growth rate ([0.28, 0.76] log d^−1^). This suggests that viral dynamics after rebound in very early treated subjects resembles acute infection dynamics.

**Table 2:**
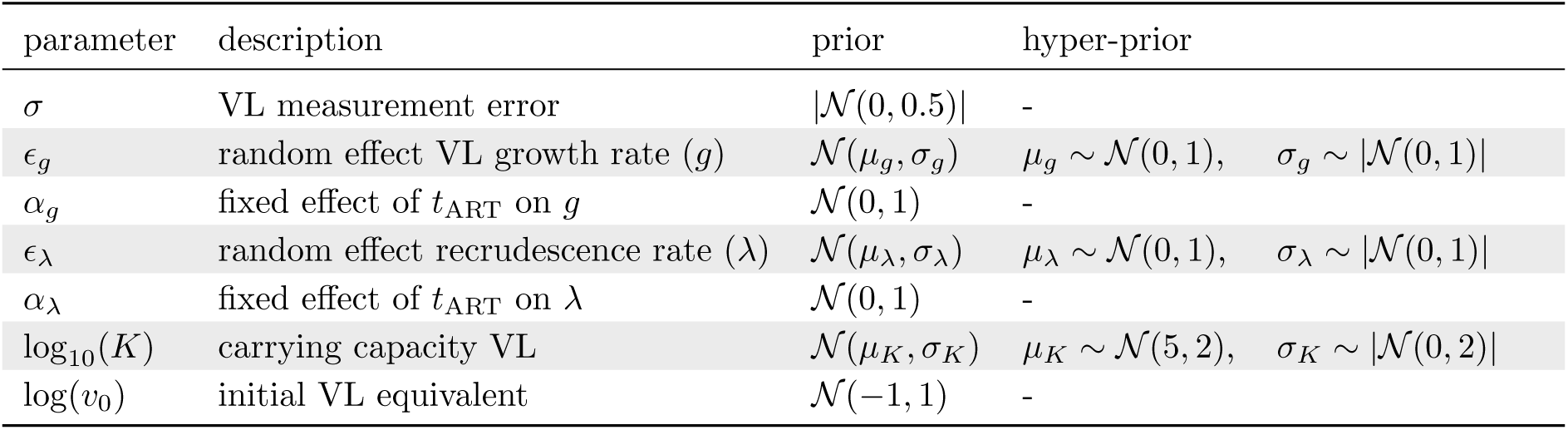
Prior distributions of the Bayesian mixed-effects model. The notation *x* ∼ *|D|* for probability distribution *D* means that *x* is positive and that *D* is truncated at zero. The normal distribution is parameterized with the mean and standard deviation. The time *t*_ART_ denotes the number of days post infection at which antiretroviral treatment was initiated.

We then used model selection theory to compare the multiple-reactivation model to the single-reactivation model (see Methods). Using the Watanabe-Akaike information criterion [WAIC; 42, and see Methods] for model comparison, we find “very strong evidence” [*sec.* 21] in favor of the multiple-reactivation model (ΔWAIC = 11.5). The superior performance of the multiple-reactivation model can be explained by two mechanisms that were mentioned above: (i) the stochasticity of secondary recrudescence events and (ii) early faster-than-exponential growth. We will now look closer into the effects of these mechanisms in the context of our SIV data set.

#### Uncertainty due to secondary recrudescence events

According to the multiple-reactivation model, successful reactivation events that follow the first event lead to faster-than-exponential growth of the VL during the early stages of rebound (Fig 1). However, these secondary reactivation events can only contribute noticeably to the viral load when the VL is still relatively low and close to the initial value *v*_0_. This most likely happens when the reactivation rate is large or the exponential growth rate is small. In order to quantify these effects, we can decompose the variance of the time-to-rebound (*τ*) as the sum of the variance of the first reactivation time (*T*_1_), and the variance due to all subsequent reactivation events. The proportion of the total variance that is due to secondary reactivation events is shown in Fig 4C for different values of the growth rate *g* and the recrudescence rate *λ*. The dark region in this heat map corresponds to a part of the model’s parameter space where it is indistinguishable from the single-reactivation model. On the contrary, the light region corresponds to parameter combinations for which most of the variance in the time-to-rebound is due to the secondary successful reactivation events. In this parameter regime the model is most relevant.

The superior performance of the multiple-reactivation model can be explained by having data from macaques with a high recrudescence rate *λ* and a small exponential growth rate *g*. Point estimates (i.e., modes of the marginal posterior distributions) of *λ* and *g* for each macaque are projected onto the heat map in Fig 4C. The macaques that were treated starting at 7, 10 and 14 days after infection fall into the parameter domain where the multiple-reactivation model is most relevant.

To further assess the effect of multiple reactivation events for each macaque, we sampled from the posterior predictive distribution of the time-to-rebound (Fig 3B, purple distributions). This distribution indicates when viral rebound is most likely to take place, given estimates for the growth rate *g*, the rate of successful reactivation *λ* and the initial viral load when exponential growth begins *v*_0_. The actual estimates for the rebound time *τ* (Fig 3A and S1, black vertical lines) correspond well with the posterior predictive distributions, as all 25 estimates of *τ* fall within the central 95% CrI of the posterior predictive distributions and 21 out of 25 estimates fall within the 50% CrI. In the model, we explicitly estimate the first recrudescence time *T*_1_ (see Methods) and hence, we can also sample from the posterior predictive distribution of *τ* conditioned on *T*_1_ (Fig 3B, green densities). These second posterior predictive distributions indicate the uncertainty in the rebound time due to secondary recrudescence events (in addition to uncertainty in the parameter estimates). Hence, by comparing the conditional (Fig 3B, green) and unconditional (Fig 3B, purple) posterior predictive distributions of *τ*, we see what effect multiple recrudescence events have on the uncertainty of the rebound time. For early treated macaques (ART *≤* 3 days post infection), most uncertainty in the rebound time comes from the first successful reactivation, as illustrated by the purple densities being much wider than the green densities. On the other hand, for the macaques treated later the subsequent reactivation events determine the rebound time distribution, as illustrated by the purple and green densities overlapping.

#### Early faster-than-exponential growth

When the recrudescence rate is large, and before the VL has become detectable, the multiple-reactivation model predicts that the VL grows faster than exponentially (Fig 1). To demonstrate the effect of this faster-than-exponential growth, we can use the regular exponential growth model to extrapolate what the first reactivation time would have been under the single-reactivation model. This time is denoted 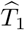, and can easily be calculated using the model’s parameters as 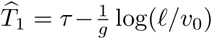. The marginal posterior densities of the first reactivation time *T*_1_ and the extrapolated initial recrudescence time 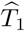 are nearly identical for the early treated macaques (Fig S2). However, for the macaques that are treated later, the extrapolated recrudescence time becomes negative (i.e. successful reactivation is predicted to occur before treatment interruption), while, according to our models, the first recrudescence time *T*_1_ has to be positive. This shows that given the estimates of *g, v*_0_, and *τ*, faster-than-exponential growth as predicted by the multiple-reactivation model is required to explain the VL data.

To identify which of the two mechanisms (uncertainty due to secondary recrudescence events or early faster-than-exponential growth) described above is the most important for explaining the difference in WAIC between the single and multiple-reactivation model, we also fit the “conditionally deterministic” multiple-reactivation model [34, and see Methods] to our SIV data set. Recall that in this model all secondary recrudescence events occur at fixed intervals. Again using WAIC, we find that our fully stochastic multiple-reactivation model fits the data better than the conditionally deterministic version, but only with limited statistical significance (ΔWAIC = 2.1 and see Table S2). As our fully stochastic model differs from the conditionally deterministic model in that it describes uncertainty in the rebound time due to secondary recrudescence events, we find that for this data set the randomness of these secondary events may be of limited importance.

### Sensitivity to parameter and model misspecification

Next, we investigated the effects of uncertainty in the initial viral load parameter (*v*_0_) and heterogeneity of the exponential viral growth rate, which can exist when the reservoir is comprised of a variety of phenotypically distinct SIV clones, on the estimates of the recrudescence rate *λ*.

#### Uncertainty in the initial viral load equivalent

The meaning of the parameter *v*_0_ is biologically ambiguous. Previously, this parameter has been described as the initial “plasma viral load equivalent” [33] and estimates of *v*_0_ are at least roughly compatible with the number of virions produced by one productively infected cell, the clearance rate of virions [38] and the blood volume of a macaque [33]. Another interpretation is linked to extinction probabilities of a recently reactivated lineage. In this case *v*_0_ is the viral load at which extinction of an exponentially growing lineage is extremely unlikely [19, 17]. The actual model is agnostic with respect to the interpretation of *v*_0_, which can be thought of as the effect size of the multiple-reactivation model. This means that *v*_0_ simply provides a measure of the effect of each independent recrudescence event, each possibly originating from a separate anatomical site [11], on the VL dynamics and time-to-rebound. However, as it is difficult to estimate both the recrudescence rate *g* and the initial viral load *v*_0_ simultaneously, we investigated the effect of a misspecified *v*_0_ on the estimate of the recrudescence rate *λ*.

To assess the bias due to misspecification of *v*_0_, we simulated large data sets (*n* = 200) with various ground-truth parameter values and fit our model to the synthetic data. For simplicity, we used an exponential growth model instead of logistic growth, and removed the random effects from the statistical model. Hence, all simulated subjects share the same parameter values. The ground-truth *v*_0_ was kept constant to 0.1 copies mL^−1^, while in the statistical model, the assumed constant value of *v*_0_ was varied from 0.02 to 0.5 copies mL^−1^. Assuming an erroneous value of *v*_0_ resulted in a sizable bias in the estimate of *λ* (Figure 5A). When *v*_0_ is assumed smaller than the ground truth value, the model requires a larger recrudescence rate in order to fit the data, and vice versa. This is especially clear when the ground-truth reactivation rate is large (*λ* = 5 d^−1^). This is as expected, because again, *v*_0_ can be interpreted as the effect size of the multiple-reactivation model, and becomes more important when secondary recrudescence events are more frequent.

**Figure 5:**
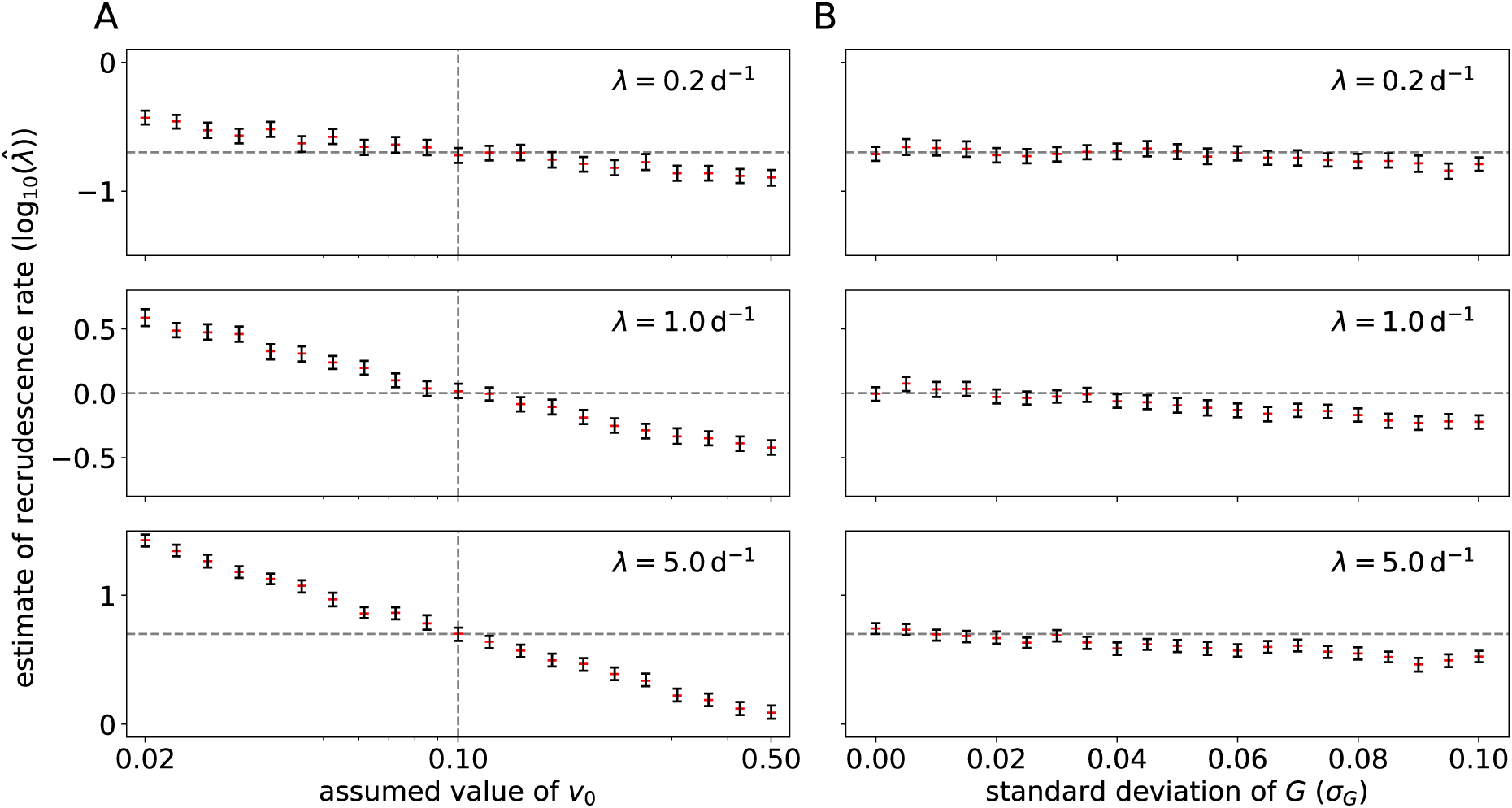
Sensitivity of the multiple-reactivation model to misspecification. (A) A mis-specified initial viral load *v*_0_ can lead to a biased estimate of the recrudescence rate *λ*. Rebound data sets (*n* = 200) were simulated by sampling from the viral load process (Eq 1) using different values of the recrudescence rate *λ* (horizontal dashed lines), and different assumed values of *v*_0_ (horizontal axis). The ground truth value of *v*_0_ equals 0.1 copies mL^−1^ (vertical dashed lines). Shown are the 95% CrIs of the estimate 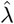 of *λ* (black bars) and the posterior medians (red). (B) Intra-host variation in the exponential growth rate of the VL can lead to a biased estimate of the recrudescence rate *λ*. Data sets of rebound time series were now simulated from a viral load process with within-host variability of the growth rate *G*_*i*_ (Eq 23, Figure S5), using different values of the recrudescence rate and the standard deviation of the viral growth rate (*σ*_*G*_, horizontal axis), ranging from 0% to 20% of the most likely growth rate *g* = 0.5.

#### Within-host heterogeneity of the exponential growth rate

Throughout the paper, we have made the strong assumption that within one subject all successfully reactivated lineages have the same exponential growth rate *g*. In natural infections, ART is only rarely started during hyper-acute infection and this means that the latent reservoir consists of a diverse archive of proviral sequences [20], probably varying in their growth rate due to intrinsic fitness costs of mutations or escape mutations from immune responses [37]. To measure the effect of this potential model misspecification, we performed a sensitivity analysis with simulated data sets as before. In this case, we had to generalize the multiple-reactivation model (Eq 1) and replace the growth rate *g* with a clone-specific random variable *G*_*i*_, representing the random growth rate of the *i*-th clone (see Eq 23 in S1 Text). Example trajectories of this generalized model are shown in Figure S5. This generalized multiple-reactivation model requires that we specify a distribution for the random growth rates *G*_*i*_. In the S1 Text we developed a simple example providing a model for an SIV reservoir in which the frequency of a clone is proportional to its fitness (see the inset of Fig S5). The most fit and abundant clone has growth rate *g* and we write *σ*_*G*_ for the standard deviation of the random growth rate *G*_*i*_.

As in the sensitivity analysis described above, we simulated large data sets (*n* = 200), varying the recrudescence rate *λ* and the standard deviation *σ*_*G*_ of the random growth rates. We then estimated the model parameters *g* and *λ* with the simplified statistical model (i.e. exponential growth instead of logistic growth and no random effects). The initial viral load *v*_0_ was kept constant to the true value. The estimated reactivation rates are shown in Fig 5B. This shows a non-zero standard deviation in the within-host growth rate introduces a bias in the estimate of *λ*. When *σ*_*G*_ is large, the estimate of the recrudescence rate is smaller than the ground-truth value. We can understand this intuitively, because clones that reactivate early might have a growth rate that is significantly smaller than the maximum growth rate *g*, which delays the time of rebound, while the observed growth rate is dominated by fitter clones that have successfully reactivated after the first clone. This effect is most pronounced when the ground-truth recrudescence rate is intermediate or large (*λ* = 1 or 5 d^−1^). Again, this is in line with expectations, because when the recrudescence rate is small, the growth rate of the total VL is mostly determined by the clone that successfully reactivates first.

In the case of the SIVmac251-infected macaques analyzed here that are treated within 2 weeks of infection, we expect that the phenotypic variation in the reactivating strains is limited and that using a constant growth rate *g* is a valid simplification. However, when a data set contains subjects that are put on ART relatively late in the acute infection, or during chronic infection, recrudescence rates estimated with the multiple-reactivation model will likely be biased downwards. We therefore investigated if our parametric rebound-time distribution can be adjusted to account for situations when *σ*_*G*_ > 0. In the S1 Text, we derive the CGF for the generalized multiple-reactivation model described above (Eq 24). In particular, the first and second cumulants can be used to derive an approximate survival function for the fraction of subjects in remission, which is in excellent agreement with simulated rebound times (Fig S6). This shows that our probabilistic methodology can be used to extend the multiple-reactivation model to account for important biological aspects as heterogeneity of the reservoir.

## Discussion

We carefully analyzed a model for SIV and HIV rebound after treatment interruption developed by Pinkevych *et al.* [34] and Hill *et al.* [19] that takes into account the potential effect of the reactivation of multiple latently infected cells on the rebound time. In doing so, we were able to derive a relatively simple statistical model that can be used for the inference of the rate of recrudescence after treatment cessation, the viral growth rate after recrudescence, and perhaps ultimately the efficacy of novel HIV treatments in delaying viral rebound. Moreover, using our mathematical formulation, the model can be compared to similar models of viral rebound in a statistically rigorous manner. We were able to find strong statistical evidence (ΔWAIC = 11.5) in favor of the multiple-reactivation model over a simple model with only one reactivation event using previously published data from treatment-interruption experiments performed in SIV-infected macaques [43, 44]. We argued that the multiple-reactivation model is most relevant for data sets that contain subjects with early viral rebound, as our SIV data set. This is often the case for human data sets as well. For example in a pooled data set of six ACTG studies [22], 6–63% of subjects showed detectable viremia within a week, and 21–74% within 2 weeks of ART cessation [10].

Our method captures the uncertainty in SIV rebound times that is due to the stochastic nature of any recrudescence events that follow the initial activation of a latently infected cell that led to remission failure. This feature is not present in the approximation derived by Pinkevych *et al.* [34]. This novel aspect slightly improves the model’s ability to describe experimental data; when we compared our fully stochastic multiple-reactivation model with the conditionally deterministic model in the context of our SIV rebound data set, we found a small ΔWAIC of 2.1 in favor of the fully stochastic model. This indicates that the most important advantage of the multiple-reactivation model is the ability to explain fast rebound due to early faster-than-exponential viral growth.

Our fully stochastic multiple-reactivation model suffers from some of the same limitations as previous renditions [33, 13, 35]. The exact biological meaning of the initial viral load parameter *v*_0_ is ambiguous, and as we have shown with our sensitivity analysis, the estimate of the recrudescence rate is biased when the value of *v*_0_ is misspecified. In our Bayesian data analysis, we resolved this issue by choosing a broad prior distribution for *v*_0_, such that uncertainty in this parameter is propagated to uncertainty in the recrudescence rate *λ*. However, the model is still sensitive to the exact location and spread of this prior distribution. In addition, we found that the multiple-reactivation model is also sensitive to within-host variation in the viral growth rate. Even though this will likely not affect our estimates, because early treatment limits the heterogeneity of the reservoir, this bias should be taken into account when the model is applied to treatment interruption experiments with later-treated subjects, in particular in most human studies.

By specifying the model in terms of the recrudescence rate *λ*, the recrudescence times *T*_*i*_, the initial viral load equivalent *v*_0_, and the exponential growth rate *g*, we have combined all complex dynamics of reactivation and the initial stochastic growth into a single abstract recrudescence event. *In vitro* experiments have pointed out that this may be an oversimplification [16]. It is likely that a reduced exponential growth rate, for instance due to a therapeutic vaccine, also influences the rate of recrudescence, because the chances of successful reactivation are dependent on the fitness of the clone, which will be influenced by the immune response. Therefore our parameters *λ* and *g* are *a priori* dependent. A possible solution would be to parameterize the model in terms of the reactivation rate instead of the recrudescence rate, and add a parameter that is determines the probability of successful reactivation. This parameter is known as the “establishment probability”, and depends on the viral dynamics in a non-trivial manner [17]. For the aims of our current analysis, the exact relation between the reactivation and recrudescence rate are not important. However, when the multiple-reactivation model is applied to novel HIV therapies that aim to (indefinitely) extend remission, it can be important to distinguish the effects of therapies that reduce viral fitness, such as therapeutic vaccination [3] or broadly neutralizing antibodies [4], and therapies that reduce the reactivation rate, such as latency reversing agents [23].

In the presented model formulation and inference, we have ignored the period of drug washout after treatment interruption. While pharmacokinetics and dynamics may be important for precisely estimating the reactivation rate, and for instance the value of *v*_0_ [33], taking a drug washout time of 0 days is a conservative assumption for the purpose of this study. Indeed, incorporating a drug washout decreases the time available for exponential growth and hence multiple reactivation events that lead to faster-than-exponential growth become more important for rapidly rebounding macaques. We verified this by repeating the analyses with a fixed drug washout period of 1 day, during which recrudescence is not allowed to occur (not shown). Compared to the single-reactivation model, the evidence in favor of the stochastic multiple-reactivation model increased (ΔWAIC = 14.3), and compared to the conditionally deterministic multiple-reactivation model results were as before (ΔWAIC = 2.2).

Based on our estimates of the effect of the ART initiation time on the recrudescence rate (*α*_*λ*_), we predict that each day that ART initiation is delayed, the recrudescence rate increases by 36%. Recently, the aforementioned genetically barcoded SIV rebound experiments [13] have been repeated with ART initiated at day 10 and 27 post infection as opposed to day 4 [32]. These barcoded experiments could in principle give a much better estimate of the recrudescence rate, because for each macaque multiple successful reactivation events can be observed by counting the frequencies of different SIVmac239M clonotypes. In the same study, the size of the reservoir was also estimated more directly by measuring cell-associated (CA) SIV DNA in peripheral blood mononuclear cells (PBMCs). Surprisingly, while the estimated size of the reservoir based on SIV CA-DNA at the time of treatment interruption is increased more than a 100-fold when ART is started at day 10 instead of day 4 post infection, the rate of successful reactivation (inferred by counting clonotype frequencies) only increases 3.6-fold, which would amount to a 25% increase per day. This rate falls within the 95% CrI of our estimate (*viz.* [18%, 62%]). When treatment was initiated even later (day 27), the frequency of CA-DNA at the time of treatment interruption appeared to plateau at the same level as the day-10 treated macaques. Surprisingly, the inferred recrudescence rate dropped to only a 2-fold increase compared to the day-4 treated macaques. This strongly suggests that our result cannot be extrapolated to ART initiation beyond hyper-early infection, making it difficult to compare these results to most human studies, because treatment almost never starts this early for human subjects. For the early treated human cohort studies that do exist [e.g. 8], the comparison between macaque and human data can be aided by the fact that macaques are challenged with a much higher infectious dose, leading to a shorter eclipse phase compared to humans.

When we consider the 25 macaques used for this study, a large qualitative difference seems to exist between the animals treated within 3 days and those animals treated after 7 days (Fig S1). This can potentially be explained by the fact that in the early-treated macaques, no SIV-specific antibody, CD4^+^, or CD8^+^ T-cell responses could be detected [43], contrary to macaques treated from day 7 onward. One could even argue that as the time of ART initiation approaches the time of infection, the viral rebound dynamics after ART interruption starts to resemble those of an acute infection (Fig 4B). Similar patterns have been found for HIV-1, where ART initiation during acute HIV infection can lead to an incomplete HIV-specific humoral immune response, as measured by diagnostic assays [12, 25]. On the other hand, patients treated during Fiebig stage I or II have been shown to develop detectable HIV-specific CD8^+^ T-cell responses [27]. Although these responses are lower in magnitude and breadth than CD8^+^ T-cell responses from untreated individuals, they show enhanced differentiation into the effector-memory T-cell phenotype, leading to a more functional CD8^+^ memory T-cell pool compared to patients for whom treatment was initiated later. The effects of early ART on the formation of immunological memory and the subsequent impact on viral rebound dynamics could be resolved by experimentally filling the gap between macaques treated at day 3 and day 7, ideally incorporating immunological assays and using a barcoded strain. In order to extrapolate beyond ART initiation within two weeks, we will likely need models that explicitly incorporate immune responses and mechanisms like CD8^+^ T-cell exhaustion [6].

Mathematical models are required to bridge the gap between experimental observations made during treatment interruption experiments and the effect induced by novel curative treatments. A more accurate mathematical model will therefore increase the precision by which we can estimate reactivation rates— and importantly the uncertainty of these estimates—and infer the efficacy of such treatments. Here we showed that with the right mathematical tools, models of rebound dynamics can easily be refined, and used to measure parameters relevant for recrudescence. As we exemplified by incorporating within-host heterogeneity of the exponential growth rate, we envisage that our framework can be extended to include many other biological aspects, such as pharmacodynamics and detailed reactivation mechanics. Hopefully, this will lead to a more accurate understanding of SIV and HIV rebound kinetics and the efficacy of novel HIV therapies.

## Methods

### Data

The collection of the data is described in detail by Whitney *et al.* [43, 44]. In short, 36 rhesus macaques were infected with 500 TCID_50_ of SIVmac251. Combination antiretroviral treatment (a cocktail of tenofovir, emtricitabine, and dolutegravir) was initiated at various times post infection (6 hours, 1, 2, 3, 7, 10, and 14 days). Treatment continued for 24 weeks, and the viral load (VL) was monitored for 16 weeks after treatment interruption, while taking weekly measurements with a limit of detection of 50 RNA copies per mL.

### The cumulants of the process *V*_***t***_

The VL is modeled by the process *V*_*t*_ given by Eq 1, where {*T*_*i*_ : *i* = 1, 2, …} are the jump times of the Poisson process *N*_*t*_, with each jump reflecting a successful reactivation event from the reservoir. The derivation of the cumulants of the process *V*_*t*_ makes use of the fact that conditioned on *N*_*t*_ = *n*, the random times {*T*_1_, …, *T*_*n*_} are independent and uniformly distributed on the interval [0, *t*] [see e.g. 28].

This simply means that if one knows that *t* days after treatment interruption exactly *n* latently infected cells successfully reactivated, then there was no *a priori* preference for when these reactivation events took place within the time window. Or course, this is only true under the assumption that successful reactivation events can be accurately modeled by a time-homogeneous Poisson process. An overview of the parameters and variables used is given in Table 1.

The cumulant-generating function (CGF) of *V*_*t*_ is defined as the logarithmic moment-generating function 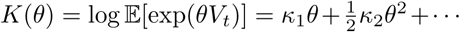*· · ·* where the first cumulant *κ*_1_ = 𝔼[*V*_*t*_] and the second cumulant *κ*_2_ = Var[*V*_*t*_]. First, assume that *N*_*t*_ = *n* so that

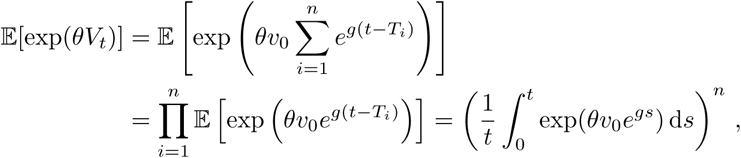

where the expectations are conditional on *N*_*t*_ = *n*. In this derivation the second equality follows from independence, and the third from the identical uniform distributions of the *T*_*i*_. Next, we drop the condition *N*_*t*_ = *n*, and use instead *N*_*t*_ ∼ Poisson(*λt*) and hence ℙ [*N*_*t*_ = *n*] = *e*^−*λt*^(*λt*)^*n*^*/n*!. Using the law of total probability, we get

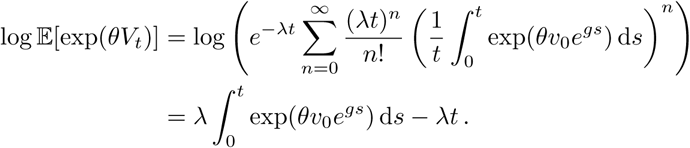

Suppose that *m* > 0. The *m*-th cumulant *κ*_*m*_ is now given by

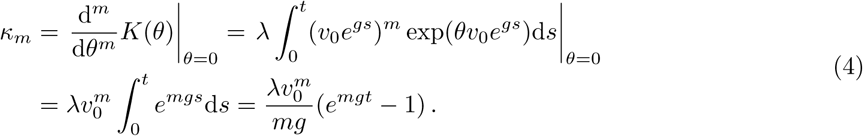

Notice that we could have used the moment generating function instead of the CGF, although calculating Eq 4 would have been more involved. A derivation of formulae for the cumulants of more general Poisson processes can be found in e.g. Privault [36].

Above we have focused on the statistics of the process *V*_*t*_ with initial condition *V*_0_ = 0. However, below we require arbitrary initial conditions *V*_0_ = *v* ≥ 0. Fortunately our results easily generalize to this situation. A VL process that starts at level *v* at time *t* = 0 can be written as *ve*^*gt*^ + *V*_*t*_ where *V*_*t*_ denotes the usual process with initial state *V*_0_ = 0. Because *ve*^*gt*^ is deterministic, the cumulant generating function of *ve*^*gt*^ + *V*_*t*_ is simply given by log 𝔼[exp(*θve*^*gt*^ + *θV*_*t*_)] = *θve*^*gt*^ + *K*(*θ*), where *K*(*θ*) is again the CGF of *V*_*t*_. Therefore, when the initial condition equals *V*_0_ = *v*, only the first cumulant (the mean) of *V*_*t*_ changes from *κ*_1_ to *ve*^*gt*^ + *κ*_1_, and all other cumulants remain unaffected.

### The derivation of the approximation 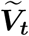

In the analysis of Pinkevych *et al.* [34], the Poisson process *N*_*t*_ is replaced by a process 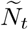 with jump times *T*_1_, *T*_2_, … that is deterministic conditioned on *T*_1_ = *t*_0_; the time of the first successful reactivation event. The subsequent recrudescence times *T*_2_, *T*_3_, … are spaced at regular intervals, with *T*_*i*+1_ − *T*_*i*_ = 1*/λ* for *i* ≥ 1. The number of successful reactivation events at time *t* > *t*_0_ is then given by 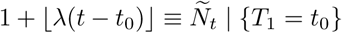, where ⌊*x*⌋ denotes the largest integer *≤ x*. Eq 2 can now be derived as follows:

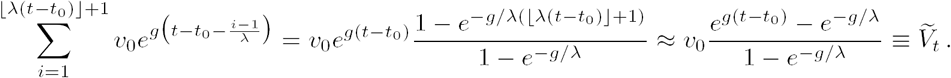

The first step follows from the identity for a geometric progression, and in the second step the approximation ⌊*λ*(*t* − *t*_0_)⌋ ≈ *λ*(*t* − *t*_0_) is used.

Another way to approximate the stochastic process *V*_*t*_, is to assume that *λ* is very large, so that latently infected cells are continuously reactivated. Each of these reactivations adds *v*_0_ SIV RNA copies mL^−1^ to the total VL. In this case it becomes feasible to use an ordinary differential equation (ODE) to describe the VL dynamics. The initial value problem (IVP) for the large-*λ* approximation 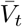 of *V*_*t*_ is given by

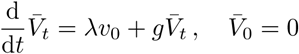

and the solution to this IVP is given by the right-hand-side of Eq 3, which was also noted by Prague *et al.* [35]. When *λ* is large, we can approximate 1 − *e*^−*g/λ*^ with *g/λ*, and *e*^−*g/λ*^ with 1 in Eq 2. This implies that 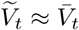 when recrudescence is fast.

### The first passage time of the limit of detection

Here, we derive a parametric probability distribution for the time to viral rebound after treatment interruption, which is our main tool for analyzing viral rebound data. Above, we have seen that the expectation of *V*_*t*_ is given by the first cumulant, 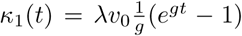, and that the variance equals 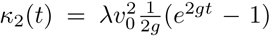. A naive way to derive an approximation for the time-to-rebound *τ* is to approximate the distribution of *V*_*t*_ with *𝒩* (*κ*_1_(*t*), *κ*_2_(*t*)), a normal distribution with mean *κ*_1_(*t*) and variance *κ*_2_(*t*), and this is essentially what we will do below. However, in the S1 Text, we will give a theoretical justification for this naive approach and use the Kramers-Moyal expansion to replace *V*_*t*_ with a transient Ornstein-Uhlenbeck (OU) process [see e.g. 41, 39].

Armed with a Gaussian approximation of the distribution of *V*_*t*_, we can derive an approximation of the distribution of the viral rebound time. Although numerical methods exist to compute the density of the true first passage time of the transient OU process *V*_*t*_ [1], here we make the assumption that the LoD *ℓ* for the VL is much larger than the initial value *v*_0_, such that we can reasonably approximate the survival function *S*(*t*) ≡ ℙ [*τ* ≥ *t*] with the cumulative density function (CDF) ℙ [*V*_*t*_ < *ℓ*] [cf. 17]. This is a valid approximation, since *V*_*t*_ grows exponentially around the relatively large LoD *ℓ*. In order to get a probability density function for *τ*, we simply differentiate the approximated survival function *S*(*t*) with respect to *t*:

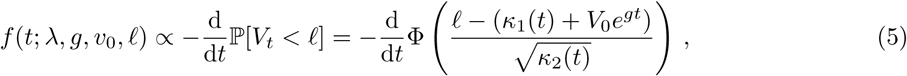

where 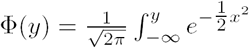 is the CDF of the standard normal distribution 𝒩(0, 1). By expanding Eq 5, we get

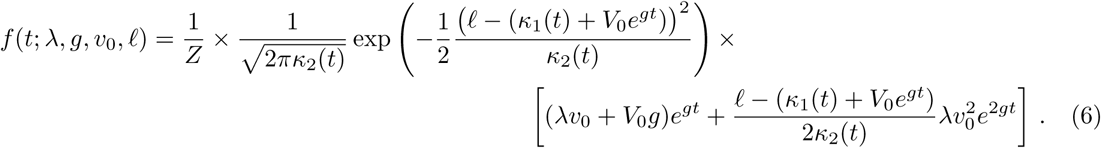

To prove that *f* is a proper probability distribution, we have to show that *f* is non-negative, and we have to find a normalizing constant *Z* for Eq 6. The reason that the right-hand-side of Eq 5 does not automatically define a proper probability density function (i.e. *Z* ≠ 1) is because the diffusion approximation of *V*_*t*_ can become negative, and declines exponentially towards −∞ with a non-zero probability. We have to condition that this non-biological event does not occur. The normalizing constant *Z* is equal to the probability of ever reaching the LoD *ℓ*:

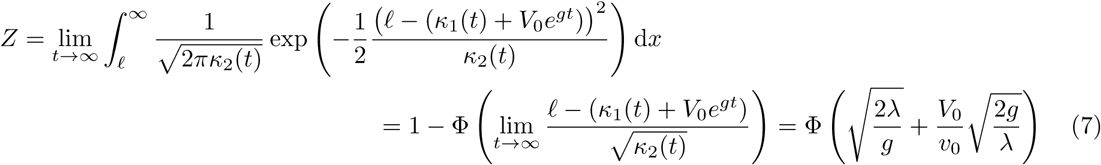

The fact that *f* is non-negative follows from a simple calculation, where we have to make the reasonable assumption the viral load at time *t* = 0 is below the limit of detection (*V*_0_ < *ℓ*):

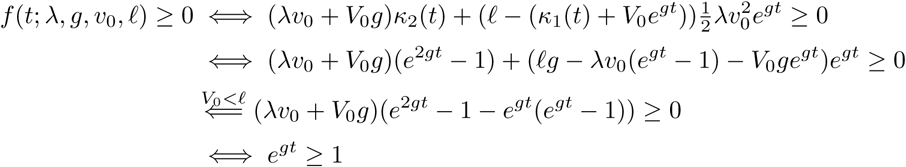

which is true for all non-negative *t*.

The expression for the rebound-time distribution *f*, given *ℓ*, allows for estimation of the parameters *λ, v*_0_ and *g* by maximization of the likelihood or other inference methods. Notice that Eq 6 and 7 somewhat simplify when we take the initial condition to be *V*_0_ = 0.

However, to justify that we can replace *V*_*t*_ with a recurrent OU process, and hence approximate its distribution with a Gaussian, we have to assume that *v*_0_ is relatively small compared to *V*_*t*_ (see S1 Text). This means that taking the initial condition *V*_0_ = 0 might be problematic. In Fig 2 we compare simulated rebound times with the approximated rebound-time distribution *f* (*t*; *λ, g, v*_0_, *ℓ*) where we have taken *V*_0_ = 0. For large *λ* the approximation and simulations are in good correspondence, but when *λ* is small we find a discrepancy. Below we solve this by taking an initial value *V*_0_ > 0.

### A mixed effects model for treatment-interruption data

Above, we derived our main tool for analyzing viral rebound data: a probability distribution for the time to viral rebound. However, in order to apply this to our SIV rebound data, we need additional statistical methodology, which we develop here. As the VL can only be observed periodically, in any treatment-interruption study the time of viral rebound *τ* is doomed to be interval-censored or right-censored. The viral dynamics after an interval-censored rebound event can be used to narrow the window in which this event occurred [35]. As the VL reaches its peak, the growth rate slows down. Therefore, using a model of pure exponential growth could easily underestimate the initial growth rate. To avoid this we use a logistic growth model with carrying capacity *K* to infer the exponential growth rate *g* and the time-to-rebound *τ* from the VL time series. Hence, at *t* days after treatment interruption, the model predicts a VL equal to

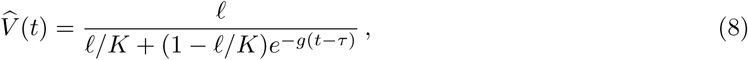

such that 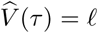. To model a proportional measurement error [24], we assume that the observed VL has a log-normal distribution around the predicted value: 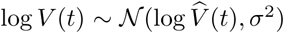. The likelihood of a left-censored observation (i.e. the VL is below the LoD) is replaced by the cumulative density of the normal distribution.

To account for the limited number of observations, we use random and mixed effects for the parameters *K, g* and *λ*. Since we know that the time of treatment initiation (*t*_ART_) is a predictor for both *λ* and *g*, we define

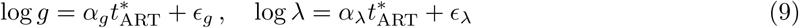

where *ϵ* _*g*_ and *ϵ*_*λ*_ are normally-distributed random effects (a standard assumption), the variable 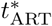 is the standardized treatment initiation time, and *α*_*g*_ and *α*_*λ*_ are fixed effects.

All we have to do now is describe a model for the parameter *τ* —the rebound time. For this we consider three different scenarios.

#### The multiple-reactivation model

In order to split the effect of the first reactivation event from subsequent events, we explicitly model the first reactivation time *T*_1_ ∼ Exponential(*λ*). The likelihood of the difference *τ* − *T*_1_ is then given by Eq 6, with initial condition *V*_0_ = *v*_0_. The parameter *v*_0_ is modeled as a fixed effect, and we chose a prior distribution around the estimates for macaques reported previously [33]. The prior distributions and hyper-parameters for all the model’s parameters are listed in Table 2. We chose broad prior distributions for all the (hyper) parameters; notice that the prior distributions are defined on a logarithmic scale.

#### The single-reactivation model

Eq 8 and 9 remain valid for the single-reactivation model and the reactivation time *T*_1_ is again assumed to be exponentially distributed with rate *λ*. However, the difference *τ* − *T*_1_ now has a Dirac-delta distribution, as it is completely determined by *g, v*_0_ and *ℓ*:

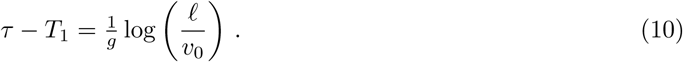

To account for this, the rebound time *τ* is no longer a free parameter in the single-reactivation model, but instead defined by Eq 10.

#### The conditionally-deterministic multiple-reactivation model

The approximation for the multiple-reactivation model that was developed by Pinkevych *et al.* [34] is deterministic after the first recrudescence event. The time between this first event and rebound can be derived using Eq 2 from solving *τ* − *t*_0_ from 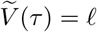, which leads to

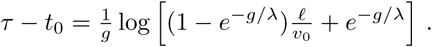

As in the case of the single-reactivation model, we let *t*_0_ = *T*_1_ ∼ Exponential(*λ*).

#### Model comparison

In order to statistically compare the three different models, we calculated the Watanabe–Akaike information criterion [WAIC; 42] as

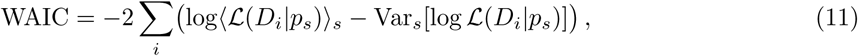

where the index *i* runs though the observations (i.e. VL measurements), and *s* runs though the Monte-Carlo samples from the posterior distribution. The function *ℒ*(*D*_*i*_*|p*_*s*_) denotes the likelihood of observation *D*_*i*_ given parameters *p*_*s*_. Moreover, we write ⟨*x*_*s*_⟩_*s*_ for the sample mean of *x* and Var_*s*_[*x*_*s*_] for the sample variance. The results of the model comparisons are listed in Table S2.

The mixed-effects model is implemented in the probabilistic programming language Stan [7]. For each model, we ran 4 independent chains of length 5000 and 1 : 20 thinning, resulting in a 1000 samples from the posterior distribution. The Gelman-Rubin statistic 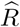 was close to 1 for all parameters, indicating good convergence of the chains. The scripts and data used for the analyses can be downloaded from https://github.com/lanl/multiple-reactivation-model.

## Acknowledgments

Portions of this work were done under the auspices of the U.S. Department of Energy under contract 89233218CNA000001 and supported by NIH grants P01-AI131365 (JBW); R01-AI028433 and R01-OD011095 (ASP); AI124377, AI126603, and AI128751 (DHB); and NSF grant DMS-1714654 (JMC). We gratefully acknowledge Garrett T. Nieddu for his technical support.

## Supporting Information

**Figure S1:**
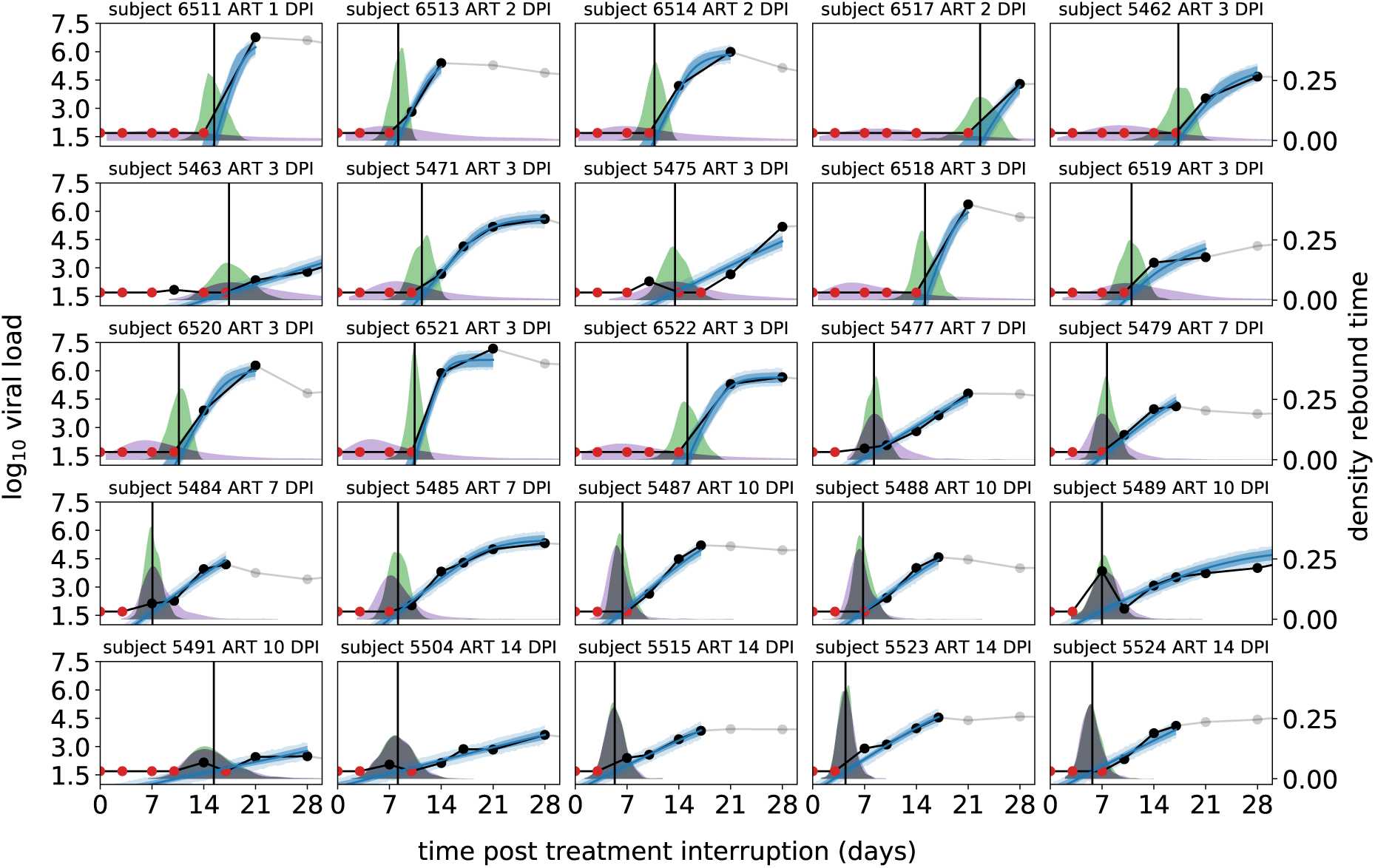
Model fits, used data points and posterior predicted rebound time distributions for all macaques. The panels (DPI: days post infection) show the VL data (black dots connected by black lines, with red dots for left-censored observations; the grey dots are ignored) taken from all 25 macaques for whom rebound was observed, and the stochastic multiple-reactivation model prediction (blue lines: posterior mean; dark blue band: 50% credibility interval (CrI), light blue band: 50% posterior predictive interval). The estimated time-to-rebound (*τ*) is given by the vertical black line. The density plots in the background indicate the posterior predictive distribution of *τ*. The green distributions are conditioned on the estimated time of the initial reactivation event, the purple distributions are unconditional.

**Table S1:** Parameter estimates and credibility intervals. Parameter estimates from the fully stochastic multi-reactivation model (“rebound” columns). and the acute infection (“acute” columns). The point estimates correspond to the mode of the marginal posterior distributions.

|  |  | *rebound* |  |  |  | *acute* |  |
| --- | --- | --- | --- | --- | --- | --- | --- |
| *population-level parameters* |  |  |  |  |  |  |  |
| symbol | unit | mode | 95% CrI |  |  | mode | 95% CrI |
| $\alpha_g$ | $\log \text{d}^{-1}$ | −0.48 | [−0.95, −0.18] | | | | |
| $\alpha_\lambda$ | $\log \text{d}^{-1}$ | 1.47 | [0.73, 2.10] | | | | |
| $\mu_g$ | $\log \text{d}^{-1}$ | −0.24 | [−0.52, 0.52] | | | 0.67 | [0.58, 0.79] |
| $\mu_\lambda$ | $\log \text{d}^{-1}$ | −0.93 | [−1.65, −0.01] | | | | |
| $\sigma_g$ | $\log \text{d}^{-1}$ | 0.63 | [0.44, 1.33] | | | 0.05 | [0.01, 0.27] |
| $\sigma_\lambda$ | $\log \text{d}^{-1}$ | 0.12 | [0.02, 0.95] | | | | |
| $v_0$ | copies mL<sup>−1</sup> | 0.36 | [0.11, 1.89] | | | | |
| $\sigma$ | log copies mL<sup>−1</sup> | 1.28 | [1.05, 1.63] | | | 0.80 | [0.51, 1.12] |
| *individual-level parameters* |  |  |  |  |  |  |  |
| subject | start ART (d) | $g$ (d<sup>−1</sup>) | 95% CrI | $\lambda$ (d<sup>−1</sup>) | 95% CrI | $g$ (d<sup>−1</sup>) | 95% CrI |
| 6511 | 1 | 2.21 | [1.24, 7.43] | 0.07 | [0.03, 0.19] |  |  |
| 6513 | 2 | 1.37 | [0.85, 4.48] | 0.10 | [0.04, 0.30] |  |  |
| 6514 | . | 1.30 | [0.77, 4.62] | 0.10 | [0.04, 0.28] |  |  |
| 6517 | . | 1.01 | [0.44, 4.60] | 0.08 | [0.04, 0.20] |  |  |
| 5462 | 3 | 0.70 | [0.42, 3.65] | 0.13 | [0.06, 0.31] |  |  |
| 5463 | . | 0.27 | [0.20, 0.43] | 0.14 | [0.07, 0.56] |  |  |
| 5471 | . | 0.82 | [0.56, 1.99] | 0.13 | [0.06, 0.35] |  |  |
| 5475 | . | 0.42 | [0.32, 0.63] | 0.14 | [0.06, 0.50] |  |  |
| 6518 | . | 1.86 | [1.09, 5.53] | 0.12 | [0.05, 0.29] |  |  |
| 6519 | . | 0.56 | [0.38, 2.88] | 0.14 | [0.06, 0.42] | 1.95 | [1.56, 2.59] |
| 6520 | . | 1.19 | [0.75, 4.05] | 0.13 | [0.06, 0.38] |  |  |
| 6521 | . | 2.36 | [1.32, 5.82] | 0.13 | [0.06, 0.32] |  |  |
| 6522 | . | 1.34 | [0.76, 5.33] | 0.13 | [0.06, 0.29] |  |  |
| 5477 | 7 | 0.56 | [0.41, 0.74] | 0.45 | [0.21, 1.96] | 1.98 | [1.73, 2.65] |
| 5479 | . | 0.67 | [0.50, 1.31] | 0.42 | [0.19, 1.78] | 1.97 | [1.64, 2.34] |
| 5484 | . | 0.65 | [0.47, 0.94] | 0.46 | [0.21, 2.45] | 2.02 | [1.81, 2.97] |
| 5485 | . | 0.65 | [0.49, 1.18] | 0.42 | [0.18, 1.62] | 1.93 | [1.57, 2.22] |
| 5487 | 10 | 0.71 | [0.54, 1.20] | 0.92 | [0.35, 4.96] | 1.90 | [1.60, 2.21] |
| 5488 | . | 0.60 | [0.46, 1.00] | 0.93 | [0.34, 5.94] | 1.96 | [1.71, 2.50] |
| 5489 | . | 0.39 | [0.26, 1.05] | 1.38 | [0.41, 12.64] | 1.91 | [1.55, 2.25] |
| 5491 | . | 0.19 | [0.09, 0.31] | 0.88 | [0.36, 5.01] | 1.93 | [1.62, 2.36] |
| 5504 | 14 | 0.23 | [0.13, 0.35] | 3.04 | [0.76, 28.52] | 2.00 | [1.80, 2.72] |
| 5515 | . | 0.43 | [0.28, 0.66] | 3.68 | [0.79, 31.65] | 1.97 | [1.71, 2.30] |
| 5523 | . | 0.56 | [0.40, 0.85] | 3.65 | [0.78, 36.13] | 1.97 | [1.67, 2.28] |
| 5524 | . | 0.45 | [0.29, 0.67] | 3.13 | [0.63, 29.46] | 1.90 | [1.62, 2.17] |

**Figure S2:**
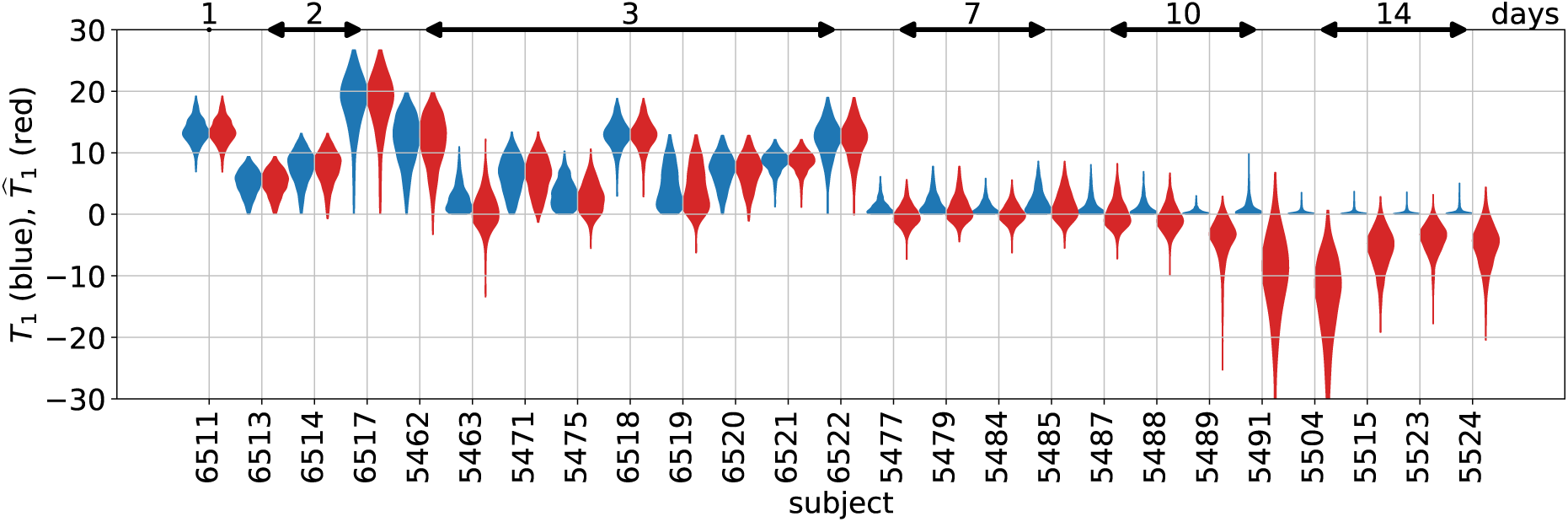
Marginal posterior densities of the first recrudescence times. Marginal densities of *T*_1_ (blue) and the extrapolated 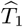 (red) for each macaque are estimated with our multiple-reactivation model. The numbers on top indicate the time of ART initiation.

**Table S2:** Watanabe-Akaike information criterion for the three models. The Watanabe-Akaike information criterion (WAIC) is averaged over 10 MCMC runs to account for Monte-Carlo error, which is indicated by the standard error of the mean (SEM). The variance 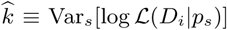 in Eq 11 can be interpreted as the effective number of parameters.

| Model | WAIC | (±SEM) | ΔWAIC | (±SEM) | $\hat{k}$ | (±SEM) |
| --- | --- | --- | --- | --- | --- | --- |
| multiple-reactivation model (MRM) | 346.3 | ($\pm 0.3$) | — | | 37.8 | ($\pm 0.2$) |
| conditionally-deterministic MRM | 348.4 | ($\pm 0.4$) | 2.1 | ($\pm 0.5$) | 38.2 | ($\pm 0.3$) |
| single-reactivation model | 357.8 | ($\pm 0.7$) | 11.5 | ($\pm 0.8$) | 42.3 | ($\pm 0.4$) |

### S1 Text

Here we derive the diffusion approximation of *V*_*t*_ by applying the Kramers-Moyal expansion to the master equation of the stochastic process *V*_*t*_. Further, we explore two other approximations of *V*_*t*_. First, we substitute the Gaussian distribution of *V*_*t*_ at time *t* with a Gamma distribution. Second, we replace the Kramers-Moyal expansion with the Wentzel–Kramers–Brillouin *ansatz*. Finally, we consider a generalization of the stochastic multiple-reactivation model that takes into account within-host variation in the exponential growth rate. We first derive the CGF, and then use the Gamma-distribution method to again derive an approximate rebound-time distribution for this generalized model.

### Theoretical justification for the Gaussian approximation of *V*_*t*_

In the main text, we derived that the stochastic process *V*_*t*_ with initial condition *V*_0_ = 0 has mean 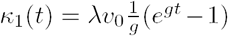 and variance 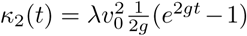. We then assumed that we could approximate the law (distribution) of *V*_*t*_ with *𝒩* (*κ*_1_(*t*), *κ*_2_(*t*)), from which we derived a probability distribution of the rebound time (see Methods). Here we will give some additional mathematical arguments to justify this approach. We will first construct a stochastic differential equation (SDE) for *V*_*t*_ with jumps given by a Poisson process with intensity *λ*. We then infer the master equation for the process *V*_*t*_, and use the Kramers-Moyal expansion to derive a Fokker-Planck equation for an approximation of *V*_*t*_. In the SDE for this approximation the Poisson process is replaced by a Brownian motion with drift *λ* and diffusion 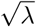. The SDE for the approximation of *V*_*t*_ can be solved explicitly, as it is the SDE for a transient Ornstein-Uhlenbeck (OU) process. For more details about these techniques, see Van Kampen [41] and Steele [39].

The stochastic process *V*_*t*_ is the solution of the SDE

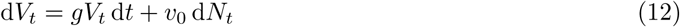

where *N*_*t*_ is a Poisson process with intensity *λ*. Let *ρ*(*t, v*) denote the distribution of *V*_*t*_. This distribution has a singular component as ℙ [*V*_*t*_ = 0*|V*_0_ = 0] = *e*^−*λt*^ *≠* 0, i.e. the VL is identically zero before the first recrudescence event. To avoid this complication, we assume that *v* ≫ *v*_0_. First, we derive the master equation for *ρ*. If *V*_*t*+*h*_ = *v*, and no reactivation has occurred in the time interval (*t, t* + *h*], then *V*_*t*_ must have been equal to *ve*^−*gh*^. The probability that no reactivation happened within this time interval is 1 − *λh*. On the other hand, if, with probability *λh*, a single reactivation did happen at time *T ∈* (*t, t* + *h*], the viral load *V*_*t*_ was equal to *ve*^−*gh*^ − *v*_0_*e*^*g*(*t*−*T*)^. Conditional on *N*_*t*+*h*_ = *N*_*t*_ + 1, the jump time *T* ∼ Uniform(*t, t* + *h*). Taking into account that probability is conserved, we get

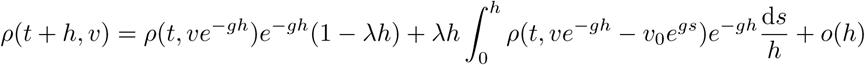

Using the mean-value theorem for integrals, we get that for some *s*^*∗*^ *∈* (0, *h*)

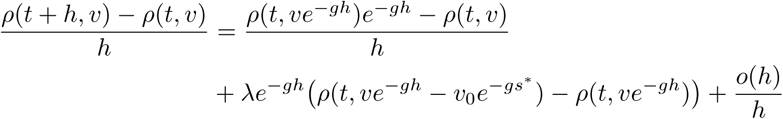

and by taking the limit *h* → 0, we find the master equation

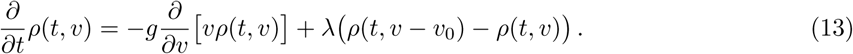

As *v*_0_ is small compared to *v*, we can use the Kramers-Moyal expansion to approximate the master equation. We first write

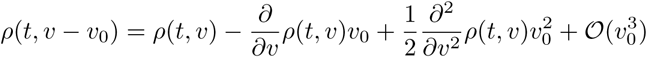

and plug this into the master equation. When we ignore terms of order 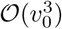, this results in the Fokker-Planck equation

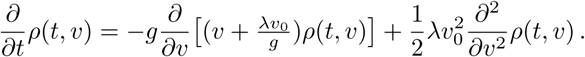

Notice that this Fokker-Planck equation corresponds to the SDE

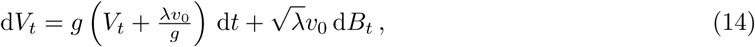

where *B*_*t*_ is a standard Brownian motion. Hence by taking the Kramers-Moyal expansion, we have replaced the Poisson process in the initial SDE (Eq 12) with 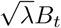, and we have added a drift term *λv*_0_d*t*. Eq 14 is up to a sign the SDE for the recurrent OU process and can be solved in a similar fashion. Let 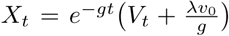 then 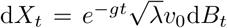, which means that 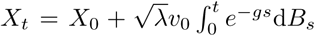.

Therefore, *X*_*t*_ is a Gaussian process with mean *X*_0_ and variance

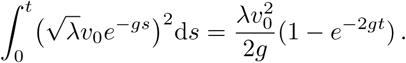

Since 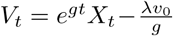, we find that *V*_*t*_ is a Gaussian process with mean 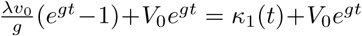 and variance 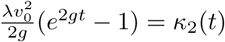.

### Alternatives to the diffusion approximation

#### Heuristically imposing a Gamma law

Above we have used the Kramers-Moyal expansion in the master equation for *V*_*t*_ to justify replacing *V*_*t*_ with a transient OU process. This led to the approximate law *V*_*t*_ ∼*𝒩* (*κ*_1_(*t*), *κ*_2_(*t*)). However, the third cumulant of the true process *V*_*t*_ is positive, and hence *V*_*t*_ is right-skewed, whereas the normal distribution is not. This suggests that we could improve the approximation of the rebound-time distribution by replacing *𝒩* (*κ*_1_(*t*), *κ*_2_(*t*)) with a right-skewed distribution. This approach is heuristic as it lacks theoretical justification.

As an example, we consider the Gamma distribution with density *v 1*→ *v*^*k*−1^*e*^−*v/η*^*η*^−*k*^Γ(*k*)^−1^, where Γ denotes the Gamma function. In order to match the first and second moments, we must have *κ*_1_ = *kη* and *κ*_2_ = *kη*^2^. We therefore get the following expressions for *k* and *η*

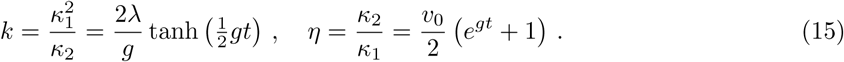

Here we have used the elementary identity *e*^2*gt*^ − 1 = (*e*^*gt*^ − 1)(*e*^*gt*^ + 1).

Write 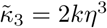 for the third cumulant of the Gamma distribution. Using our expression for *η*, we find

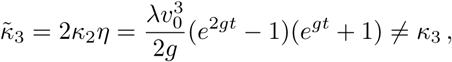

and therefore the third cumulants of *V*_*t*_ and the matched Gamma distribution do not coincide. However, the relative difference between the two cumulants *κ*_3_ and 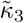 is bounded, as for all *t* ≥ 0 we have 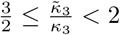. In order to see this, we notice that

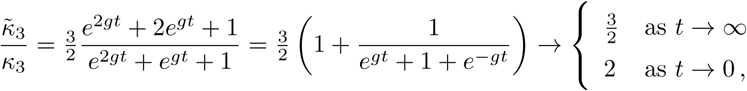

and that *e*^*gt*^ + 1 + *e*^−*gt*^ is a non-decreasing function of *t*.

Following the same steps as with the Gaussian case, we get the following survival function *S*(*t*) for the rebound time

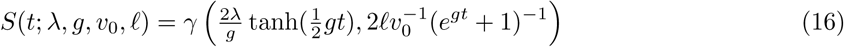

where 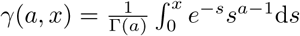 d*s* denotes the regularized incomplete Gamma function.

To prove that *S* is a proper survival function, we have to show that *S* is a monotonically non-increasing function of *t*. The derivative of *S* is equal to

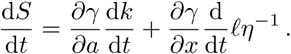

As *η* and *k* are monotonically increasing functions of *t* (see Eq 15), and *γ* is a monotonically non-decreasing function of *x*, we only have to verify that *γ* as a function of *a* is monotonically non-increasing. Using a change of variables *s* = *ux*, and splitting the Gamma function into the sum of two integrals, we get the following expression for the regularized incomplete Gamma function

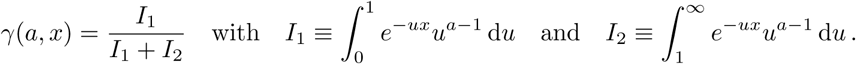

The integrand of *I*_1_ is a monotonically non-increasing function of *a*, because the integration variable *u ∈* [0, 1]. Conversely, the integrand of *I*_2_ is a monotonically non-decreasing function of *a*. Hence, *γ* monotonically decreases as a function of *a*. This shows that *S* is indeed monotonically non-increasing.

In Fig S3, we compare the rebound-time distribution corresponding to survival function (Eq 16) with simulations, using three different recrudescence rates *λ*. Comparing Fig 2 and S3 shows that the diffusion approximation and the Gamma-based approximation perform equally well for *λ* = 5 d^−1^, and 1 d^−1^. When successful reactivation events are rare (*λ* = 0.2 d^−1^), the approximation based on the Gamma law outperforms the diffusion approximation. However, the Gamma-based approximation is still unable to capture the exponential tail of the time-to-rebound distribution that is visible in the simulations with a small recrudescence rate.

The probability-density function of the time-to-rebound distribution is equal to 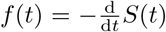. Using e.g. Mathematica [45], it is possible to obtain an expression for *f* in terms of a variety of special functions, which are not available in many other software packages. However, if the data of interest consists solely of interval- and right-censored rebound times and subsequent VL observations are not used to estimate the exact instance that the VL became observable, the density *f* is not required and the survival function *S* can be used directly [cf. 10] to calculate the likelihood of the data.

#### The WKB approximation of the master equation

In addition to the heuristic attempt to improve the approximation of the rebound-time distribution, we here explore a more advanced approach in which we replace the Kramers-Moyal expansion of the master equation (Eq 13) with the Wentzel–Kramers–Brillouin (WKB) *ansatz*. For details about this technique, we refer to e.g. Friedlin *et al.* [14]. Assuming that *v*_0_ is a small parameter, the WKB *ansatz* suggests that we write 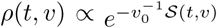 for some function 𝒮 (not to be confused with the survival function *S*). When we substitute 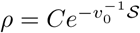 in the master equation, and divide everything by 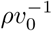, we get

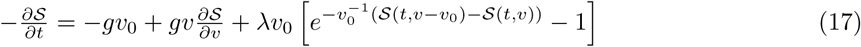

We now take a first-order Taylor expansion of 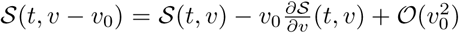 around *v* and when we ignore terms of order 𝒪(*v*_0_), we can write

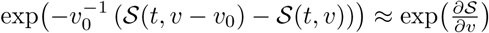

Again ignoring terms of order 𝒪(*v*_0_), Eq 17 simplifies to

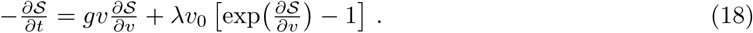

Here we have to assume that *λ*^−1^ = 𝒪(*v*_0_) as *v*_0_ → 0, but below we will see that our results hold for small *λ* as well. Hence, we have replaced the master equation for *V*_*t*_, which is both a functional and partial differential equation, with the first-order non-linear PDE in Eq 18, which can be solved with the method of characteristics. Eq 18 has the form of a Hamilton-Jacobi equation 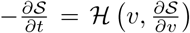 with Hamiltonian *ℋ*(*v, p*) = *gvp* + *λv*_0_(*e*^*p*^ − 1) and we find the canonical equations [see e.g. 15]

**Figure S3:**
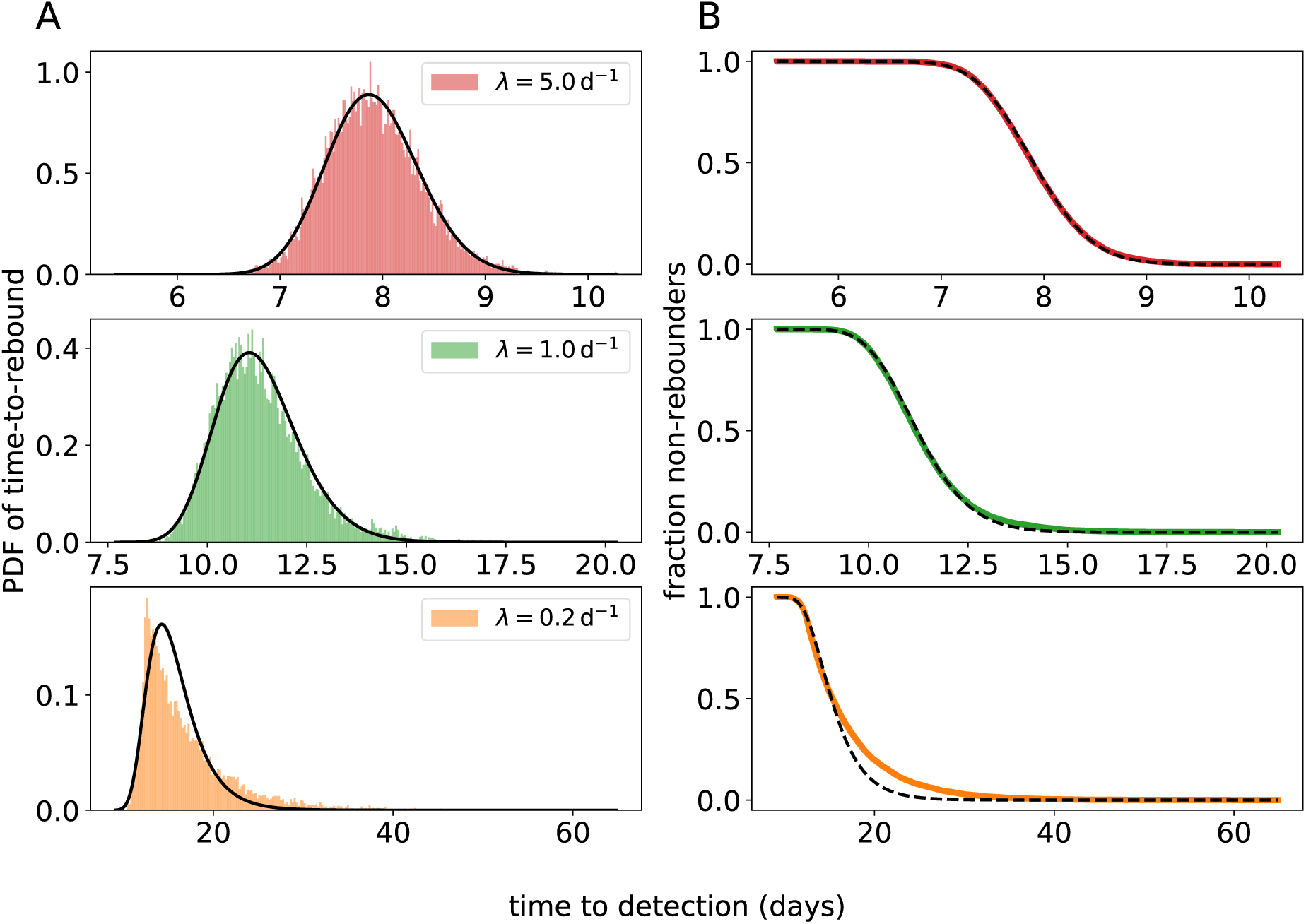
Comparison between simulated rebound times and an alternative approximation for the time-to-rebound distribution. In this case, the law of *V*_*t*_ is approximated with a Gamma distribution with mean *κ*_1_(*t*) and variance *κ*_2_(*t*). The simulated empirical distributions are shown in color, and our approximation is shown in black. The predicted PDF (A) is calculated with numerical differentiation. (B) The survival function (i.e. the fraction of subjects *S*(*t*) that do not have a detectable VL at time *t*) is defined by Eq 16. For the top, middle, and bottom panels different values of *λ* are used (*λ* = 5 d^−1^, 1 d^−1^, and 0.2 d^−1^ respectively). Notice the different time scale on the *x*-axes. For the remaining parameters, we used the values: *g* = 0.5 d^−1^, *v*_0_ = 0.1 copies mL^−1^, LoD *R* = 50 copies mL^−1^.

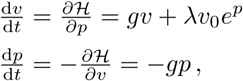

which can be solved explicitly. First, we find that *p*(*t*) = *p*_0_*e*^−*gt*^, and we get a first order ODE for *v* with time-dependent parameters and initial condition *v*(0) = 0. This ODE has solution

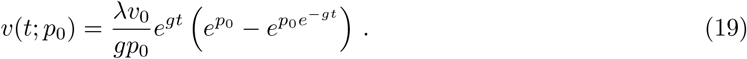

If we take the limit *p*_0_ → 0 in Eq 19, we get 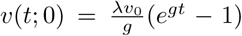, which is the trajectory of the expectation of *V*_*t*_. A solution of the PDE in Eq 18 can now we derived by integrating the Lagrangian associated with *ℋ* along the characteristic paths *v*(*t*; *p*_0_). This Lagrangian *ℒ* is given by

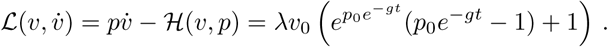

Here we write 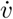 to denote the time-derivative of *v*. Hamilton’s principal function is then given by the integral of the Lagrangian along a characteristic path {*v*(*s*; *p*_0_) : *s ∈* [0, *t*]}, i.e.

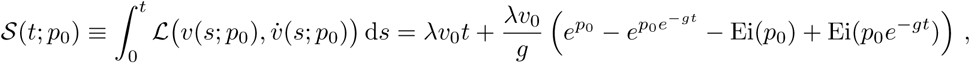

where 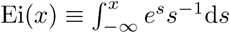 d*s* denotes the exponential integral. In order to find a solution 𝒮(*t, x*) of Eq 18, we have to find a *p*_0_ = *p*_0_(*t, x*) such that *v*(*t, p*_0_(*t, x*)) = *x*. Then 𝒮(*t, x*) = 𝒮(*t*; *p*_0_(*t, x*)).

It turns out that as an approximation of the rebound time, we can simply take

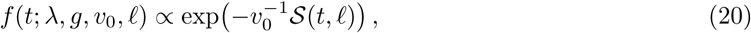

which can be made precise using the theory of large deviations [14]. In Fig S4, we have compared the rebound-time distribution derived using the WKB approximation with simulated rebound times. Comparing this with Fig 2 and S3, we find a significant improvement in the accuracy when *λ* is small. Hence, the WKB approximation is much better at describing the exponential tail of the rebound-time distribution that is due to the exponential waiting time of the first successful reactivation.

However, in order to apply this method, we have to solve the equation *v*(*t*; *p*_0_) = *ℓ* for *p*_0_, and find a constant that normalizes *f* in Eq 20. Both of these problems have to be solved numerically, which makes the method difficult to implement in a parameter-inference framework. We can somewhat simplify the equations by taking two limits. First, the process *V*_*t*_ is nearly deterministic above the detection limit. This is reflected by the fact that the Lagrangian vanishes as *t* becomes large. So instead of integrating the Lagrangian from 0 to *t* (assuming that *v*(*t*; *p*_0_) = *ℓ*), we might as well integrate from 0 to ∞, as the contribution from the interval (*t*, ∞) is negligible. In this case, Hamilton’s principal function is given by

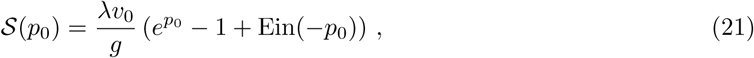

where the function 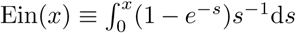 can be expressed in terms of other exponential integrals.

**Figure S4:**
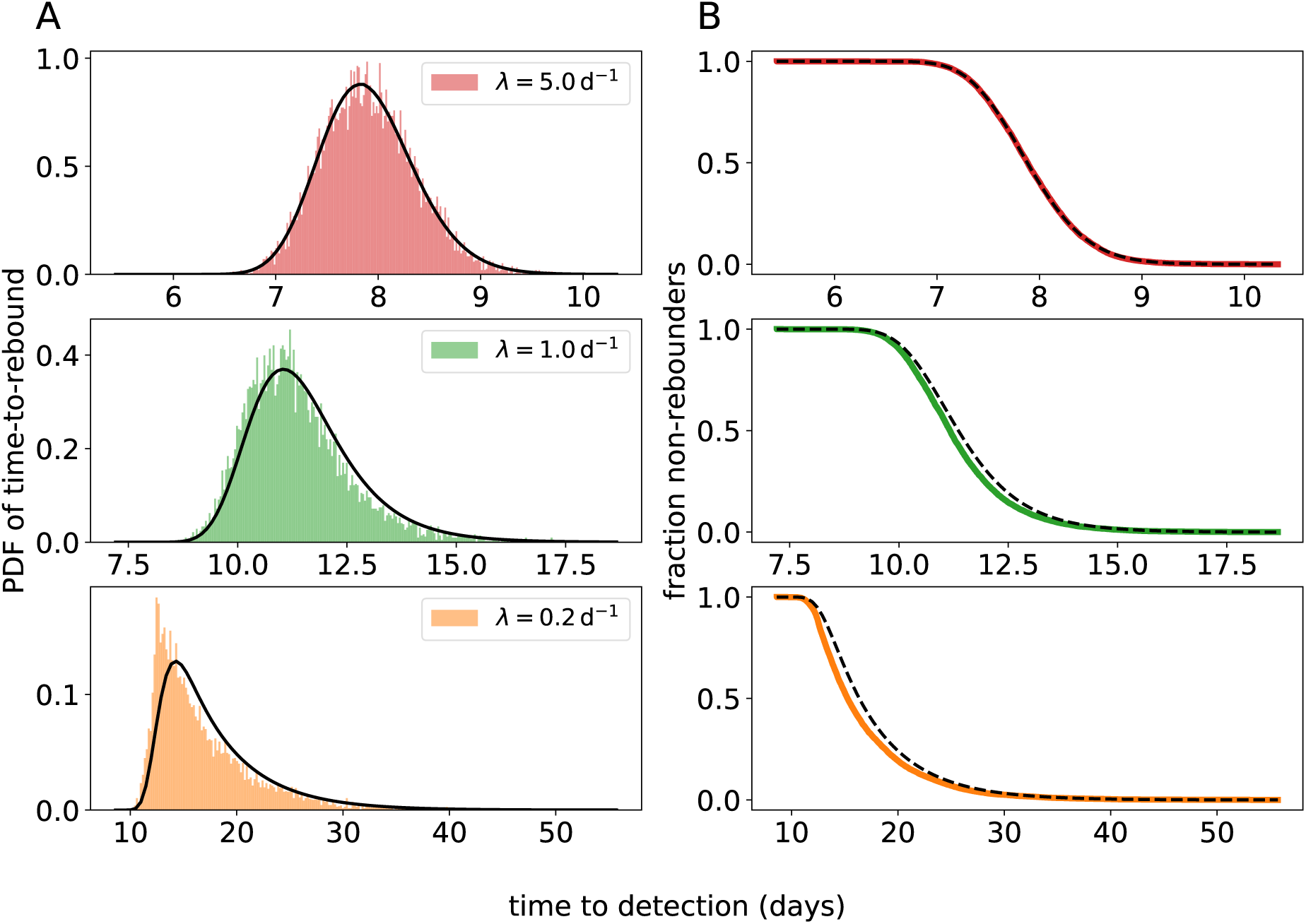
Comparison between simulated rebound times and an alternative approximation for the time-to-rebound distribution. In this case, the master equation is approximated using the WKB *ansatz*. The simulated empirical distributions are shown in color, and our approximation is shown in black. (A) The probability density function (PDF; defined by Eq 22 and Eq 21). (B) The survival function (i.e. the fraction of subjects *S*(*t*) that do not have a detectable VL at time *t*) is calculated with numerical integration. For the top, middle, and bottom panels different values of *λ* are used (*λ* = 5 d^−1^, 1 d^−1^, and 0.2 d^−1^ respectively). Notice the different time scale on the *x*-axes. For the remaining parameters, we used the values: *g* = 0.5 d^−1^, *v*_0_ = 0.1 copies mL^−1^, LoD *R* = 50 copies mL^−1^.

Second, we can make use of an asymptotic symmetry which is again due to the near determinism as *V*_*t*_ becomes large. Let *L* > *ℓ* be some VL level much larger than the LoD *ℓ*. As *V*_*t*_ grows exponentially, it takes about 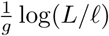 days to reach level *L* starting at LoD *ℓ*. This means that the parameter *p*_0_ that solves *ℓ* = *v*(*t*; *p*_0_) must be nearly identical to the solution of 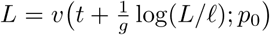. The latter equation can be re-arranged as

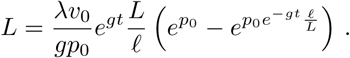

After dividing by both sides of the equation by *L*, we can take the limit *L* → ∞, and we get the following equation for *p*_0_

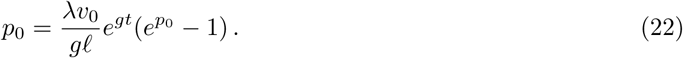

This equation can be solved in terms of the Lambert *W* function. We used Eq 22 together with Eq 21 to plot the curves in Fig S4. Despite these simplifications, using this method for inference would still be difficult due to the unknown normalizing constant in Eq 20.

#### Incorporating within-host variation in the exponential growth rate

In the models described above, we have assumed that the exponential growth rate *g* is constant within a host. Here we generalize the model so that we can incorporate variation in this growth rate. We assume again that recrudescence happens according to a Poisson process at constant rate *λ*. At each recrudescence time *T*_*i*_, a realization of the random variable *G*_*i*_ is sampled, which determines the growth rate of the *i*-th successfully reactivating clone. For mathematical tractability, we have to assume that the *G*_*i*_ are independent from each other and from *T*_*i*_ and identically distributed. In reality, this is not necessarily true, as the growth rate is related to viral fitness and clones with a higher fitness are more likely to reactivate successfully. The viral load process *V*_*t*_ at time *t* after treatment interruption is now given by

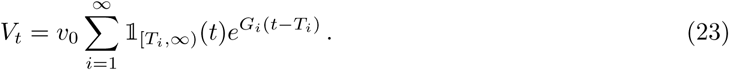

Example realizations of this process are shown in Fig S5. As before, we can derive the cumulant-generating function *K*(*θ*) = log 𝔼 [exp(*θV*_*t*_)], but now we have to take into account that the growth rates *G*_*i*_ are random variables. We first condition on *N*_*t*_ = *n* as before, and get

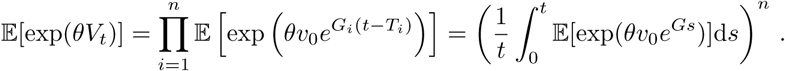

Here the expectations are conditional on *N*_*t*_ = *n* and *G* is identically distributed as any one of the *G*_*i*_. Now, we sum over all possible *n* and take the logarithm to get

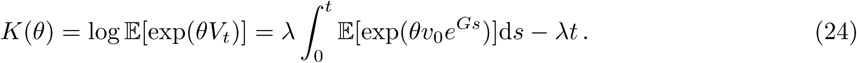

**Figure S5:**
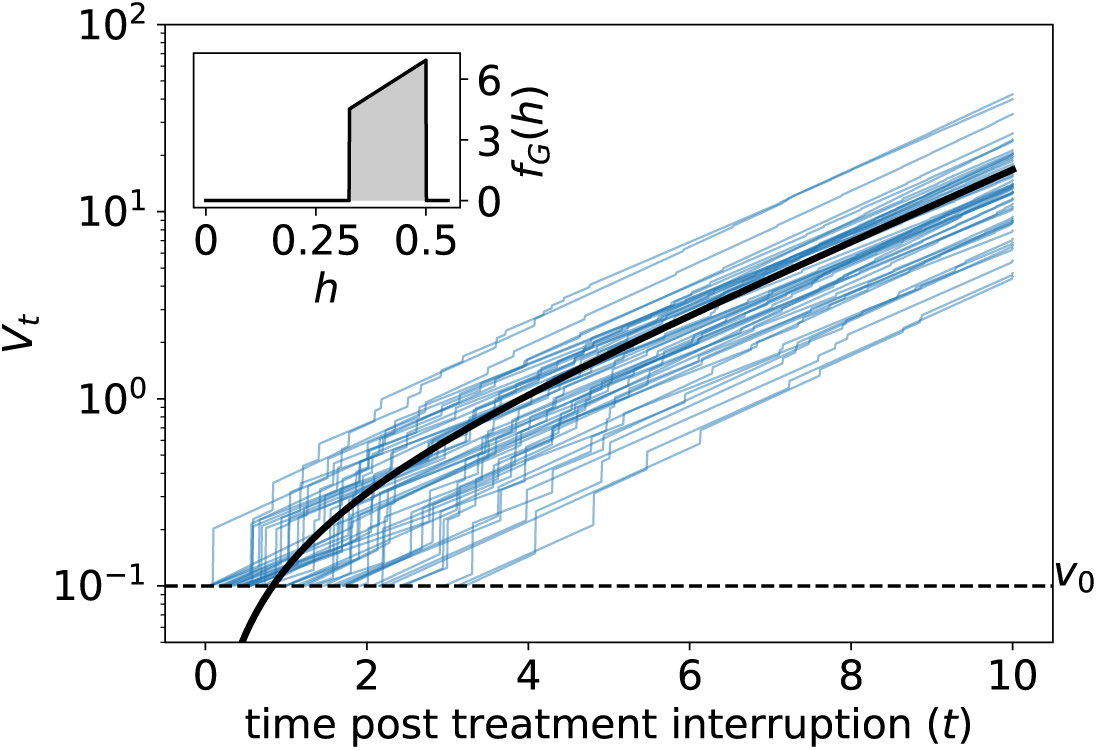
Example realizations (in blue) of the viral load process *V*_*t*_ given by Eq 23. The black curve shows the expected value 𝔼 [*V*_*t*_] = *κ*_1_ (Eq 26). The inset shows the probability density function of the random growth rate *G*_*i*_. The used parameters values are *g* = 0.5 d^−1^, *σ*_*G*_ = 0.05 d^−1^ (corresponding to *u ≈* 0.175), *v*_0_ = 0.1 copies mL^−1^, and *λ* = 1 d^−1^.

Notice that we now require that the moment generating function of *v*_0_ exp(*Gs*) exists, which is true when e.g. *G* is bounded, but not the case for arbitrary distributions of *G*. Now we can again extract the first and second cumulant by evaluating the first and second derivative of *K*(*θ*) at *θ* = 0:

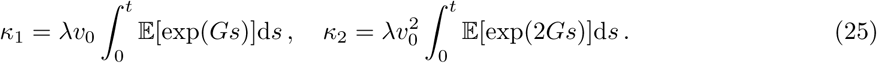

Here we require that the distribution of *G* is well-behaved enough such that we can interchange differentiation and taking the expectation. Again, this is true when we make the biologically plausible assumption that *G* is bounded.

To proceed from here, we have to choose a probability distribution for the growth rate *G*. As an example, we choose a convenient distribution that results in simple elementary expressions for *κ*_1_ and *κ*_2_. We hypothesize that clones with a higher growth rate (fitness) constitute a larger part of the reservoir, for instance because they could have been more common during acute infection. The most common clone in the reservoir has the growth rate *g*, and all other clones are less fit and have growth rates *h* in the interval [*g* − *u, g*], with likelihood proportional to *h*. Hence, the distribution of *G* is given by the PDF 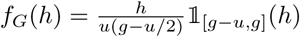 (see the inset of Fig S5). The variance of *G* is equal to 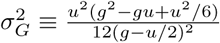 and can be adjusted by choosing the width *u* of the support of *G*.

With this choice for the distribution of *G*, we get

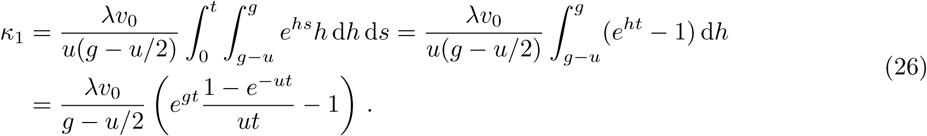

Notice that the factor *h* in the first integrand ensures that we get an elementary expression for *κ*_1_. Similarly, we get

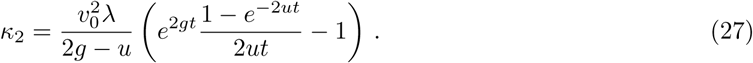

Now that we have expressions for the mean (*κ*_1_) and variance (*κ*_2_) of *V*_*t*_, we can again construct an approximate probability density function of the rebound time *τ* by approximating the distribution of *V*_*t*_ with a convenient probability distribution that has the same mean and variance. In this case, we can not take the normal distribution, as the *z*-score 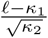 of the LoD *ℓ* is not a monotone, decreasing function of *t*. However, we can still use the our heuristic Gamma law instead of a normal distribution, with parameters 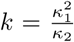 and 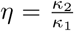. Although we could not mathematically prove that this method resulted in a well-defined survival function and PDF, we verified numerically that for biologically plausible parameters and time windows the survival function is monotonically non-increasing, and that the PDF is non-negative. The resulting PDF and survival function are compared to simulated rebound times in Fig S6. In the same figure, we have repeated the approximate rebound time distributions derived from the model with a fixed growth rate *g* (Fig S6, gray curves). This shows clearly that viral rebound is delayed in the case of a variable growth rate. This is to be expected, because the first clones that reactivate might have a smaller exponential growth rate (between *g* − *u* and *g*), and take longer to reach the limit of detection. Eventually, a clone with a growth rate close to *g* will successfully reactivate.

**Figure S6:**
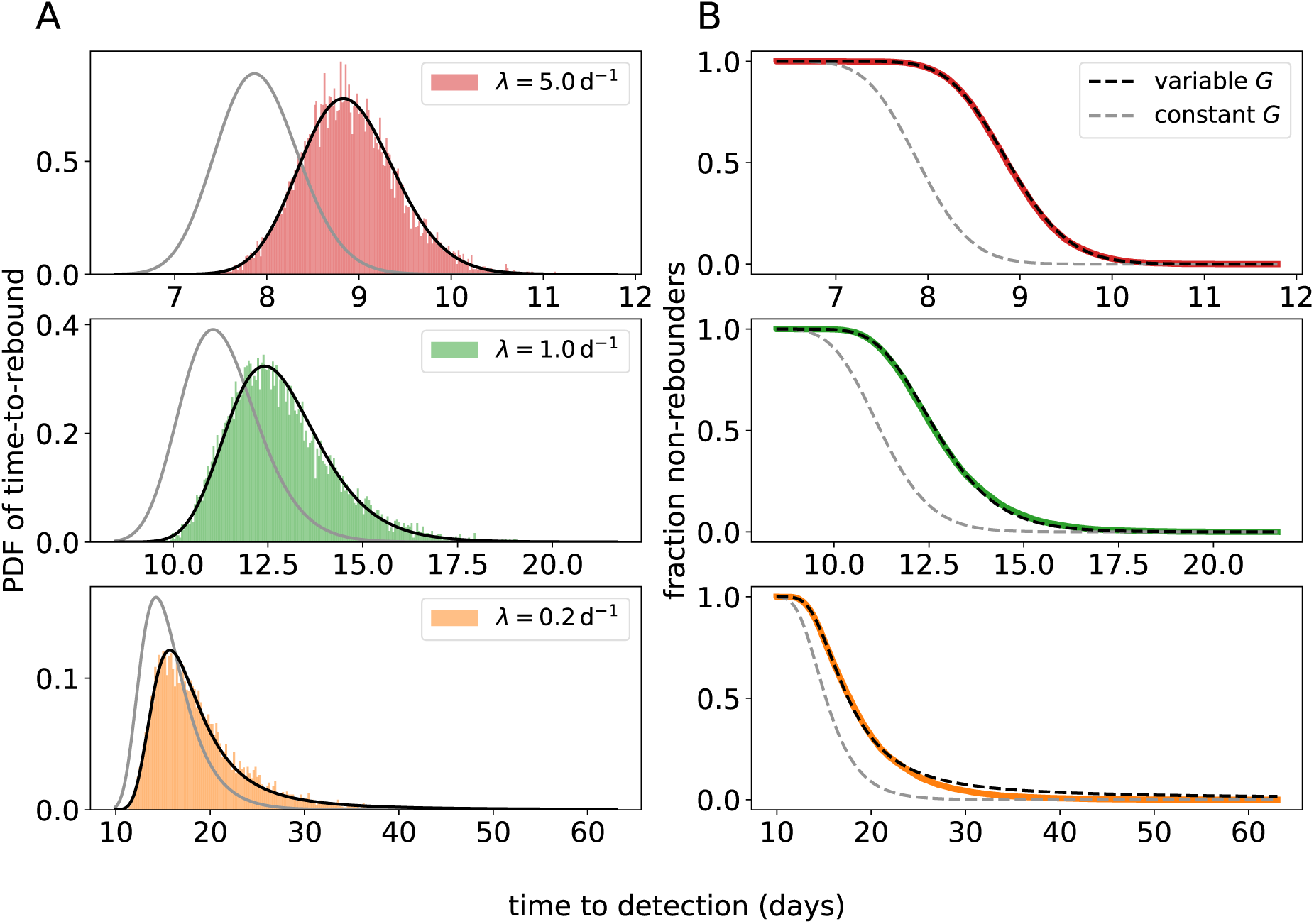
Comparison between simulated rebound times and an approximation for the time-to-rebound distribution. This model allows for variation in the exponential growth rate. The law of *V*_*t*_ is approximated with a Gamma distribution with mean *κ*_1_ (Eq 26) and variance *κ*_2_ (Eq 27). The simulated empirical distributions are shown in color, and our approximation is shown in black. The predicted PDF (A) is calculated with numerical differentiation. (B) The survival function (i.e. the fraction of subjects *S*(*t*) that do not have a detectable VL at time *t*) is defined as *S*(*t*) = *γ*(*k, ℓ/η*) with *γ* the regularized incomplete Gamma function with parameters *η* = *κ*_2_*/κ*_1_ and 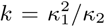. For the top, middle, and bottom panels different values of *λ* are used (*λ* = 5 d^−1^, 1 d^−1^, and 0.2 d^−1^ respectively). Notice the different time scale on the *x*-axes. For the remaining parameters, we used the values: *g* = 0.5 d^−1^, *σ*_*G*_ = 0.05 d^−1^ (corresponding to *u ≈* 0.175), *v*_0_ = 0.1 copies mL^−1^, LoD *R* = 50 copies mL^−1^. The gray curves correspond to the approximate rebound time distribution with a constant growth rate (*G* ≡ *g*) and are identical to the black curves in Fig S3.

## Notes

### Competing Interest Statement

The authors have declared no competing interest.

